# Rhythmicity of photoreceptor outer segment phagocytosis differs between the cone subtypes in the larval zebrafish

**DOI:** 10.1101/2025.01.10.632332

**Authors:** Jenni Partinen, Noora Nevala, Sanni Erämies, Teemu Ihalainen, Soile Nymark

## Abstract

Phagocytosis of retinal rod and cone outer segment (OS) tips by the retinal pigment epithelium (RPE) occurs daily to prevent accumulation of harmful compounds in the photoreceptors. Rhythmic bursts of phagocytosis, seen as increased levels of phagocytosed OS particles in the RPE, are known to appear once or twice a day depending on the animal species. However, differences in the rhythmicity of phagocytosis between distinct photoreceptor types are not well understood. Here, we show that phagocytosis of cone subtype OSs does not have identical rhythmic profiles in young zebrafish larvae. We investigated this by immunolabelling histological sections from the eyes of larvae that were collected at seven different time points throughout a 24 h circadian cycle. Internalized OS particles were then quantified from confocal images. The results revealed that OSs of all cone subtypes are phagocytosed continuously at some levels in young zebrafish. Interestingly, we observed a significant increase in the OS phagosome numbers from UV and blue cones at two time points, whereas green and red cones were phagocytosed more evenly throughout the day. We also investigated whether this rhythmicity is regulated by the external light by keeping the larvae in constant darkness before sample preparation. We found that complete darkness condition dampened the peaks in OS phagosome numbers from UV and blue cones indicating that the rhythmicity is primarily driven by the external light rather than the intrinsic circadian clocks in young larval zebrafish. Our findings provide new understanding on the rhythmicity of cone OS phagocytosis and its regulation.

## 1. Introduction

Retinal photoreceptors, rods and cones, converse light information into electrical signals that are eventually conveyed to the brain. Rods function primarily at low-light levels, whereas in most species, cones are active in bright light and are responsible for high spatial acuity and color vision. The first molecular players in the visual perception are the light-absorbing opsin proteins that are stored in the outer segments (OSs) of the photoreceptors and are specific to each photoreceptor type (Terakita 2005). Retaining their light sensitivity and overall wellbeing depends on the retinal pigment epithelium (RPE) that forms a tight interlocked structure with the photoreceptor OSs via the apical microvilli (Strauss 2005). OSs of retinal photoreceptors are continuously exposed to high-energy light from the environment, and thus, they are prone to light-induced damages. To prevent accumulation of harmful photo-oxidative molecular compounds in the photoreceptors, the OS tips are phagocytosed by the RPE while new OS membrane is continuously generated at the OS base (Young 1967; Young and Bok 1969; Lakkaraju et al. 2020). In the phagocytosis process, the most distal and aged parts of the OSs, where the phototoxic waste material has been stored are taken up by the RPE as small particles (Lakkaraju et al. 2020). After particle internalization, formed OS phagosomes are moved from the apical to the basal region inside the RPE, and simultaneously, OS phagosome maturation occurs as they interact with acidic endosomes and lysosomes (Herman and Steinberg 1982a, b; Lakkaraju et al. 2020). Eventually, the OS phagosomes are degraded in the RPE. Current knowledge on the process is mainly based on rods, but little is known about the regulation of cone OS phagocytosis.

OS tip phagocytosis is a daily occurring highly cyclic process that has one or two clear activity peaks, defined as a significant increase in phagocytosed OS particles, during a 24 hour (h) diurnal cycle depending on the animal species and their rod/cone ratio (Vargas and Finnemann 2022; Moran et al. 2022b). For instance, in rod-dominant nocturnal animals, such as rat *(Rattus norvegicus)* and northern leopard frog (*Rana pipens*), the burst of phagocytosis occurs once per day after light onset in the morning ( Basinger et al., 1976; LaVail, 1976). On the contrary, for diurnal cone-dominant animals, such as Sudanian grass rat (*Arvicanthis ansorgei*) and zebrafish (*Danio rerio*), two peaks have been observed (Bobu & Hicks, 2009; Lewis et al., 2018, Figure 3A). The peaks of OS phagocytosis in cone-dominant animals are also often associated with the change in light condition. For example, in 14-16 *days post fertilization (dpf)* zebrafish, the first peak appears 4 h after light onset in the morning, whereas the second peak emerges 3 h after light offset in the evening (Lewis et al. 2018; Moran et al. 2022a). Additionally, the dependence of the process on the endogenous circadian clocks, the intrinsic timing mechanisms driving the day-night cycles of organism’s physiology (Mohawk et al. 2012; Finger and Kramer 2021), has been investigated. In many mammals, such as mice and rats, the rhythmic nature of the process remains in constant darkness, demonstrating circadian regulation (LaVail 1980; Grace et al. 1999). However, in some vertebrates, such as frogs and goldfish (*Carassius auratus*), the peaks of OS phagosomes disappear when the ambient light is removed (Basinger and Hollyfield 1980, Bassi et al., 1990), suggesting that the rhythm is driven primarily by the external light-dark cycle. Similar preliminary evidence is received from 16 *dpf* zebrafish; the peaks disappear in constant darkness (Moran et al. 2022a). However, it is worth noting that, the Moran et al. data is based on three time points only, and thus, do not take into account the possibility that removing the external light might shift the peak time of phagocytosis towards earlier or later hours of the day.

Most of the studies regarding the OS phagocytosis are conducted in animals with mature retinas (LaVail 1980; Bobu and Hicks 2009; Lewis et al. 2018; Milićević et al. 2021), leaving characteristics of the process in young animals with developing retinas elusive. In fact, 14-16 *dpf* old zebrafish were previously used to reveal the daily peaks of OS phagocytosis (Lewis et al. 2018; Moran et al. 2022a), but the rhythmicity has neither been studied in younger nor older zebrafish, where the retina is at different developmental stages. When the zebrafish retina has reached its anatomical maturity at 20 *dpf*, retinal cross-sections and flat mounts reveal multilayer organization of OSs of different photoreceptor types: four types of cones (ultraviolet (UV), blue, green and red) and one type of rods (Branchek and Bremiller 1984; Allison et al. 2010). Instead, in larval zebrafish, the photoreceptors are approximately in the same layer up to 8 *dpf* (Branchek and Bremiller 1984; Allison et al. 2010). The OSs of different photoreceptor types can be distinguished from 4 *dpf* onwards (Crespo and Knust 2018), red/green double cones start to appear at 12 *dpf* and all cone subtypes reach their full adult dimensions at 15 *dpf* (Branchek and Bremiller 1984). Rods become mature even later, at 20 *dpf* (Branchek and Bremiller 1984; Bilotta et al. 2001; Morris and Fadool 2005). As zebrafish develops, also the configuration of the photoreceptor OSs evolve continuously towards more organized conformation. Simultaneously, the physical interaction and the interactive processes between the photoreceptors and the RPE become more evident as the animal grows. An example of these interactive processes are retinomotor movements, seen as changes in photoreceptor OS positioning in relation to RPE tissue due to the light and circadian rhythms in teleost fishes (Hodel et al., 2006; Menger et al., 2005, Figure 1A). These movements of rods and cones are converse: in the light, rods elongate deeper in the RPE while cones contract and in dark conditions, rods contract while cones elongate (Menger et al., 2005, Figure 1A). In contrast to adults, retinomotor movements of rods and cones are weak in the larval zebrafish retina under 28 *dpf* (Hodel et al. 2006).

**Figure 1.**
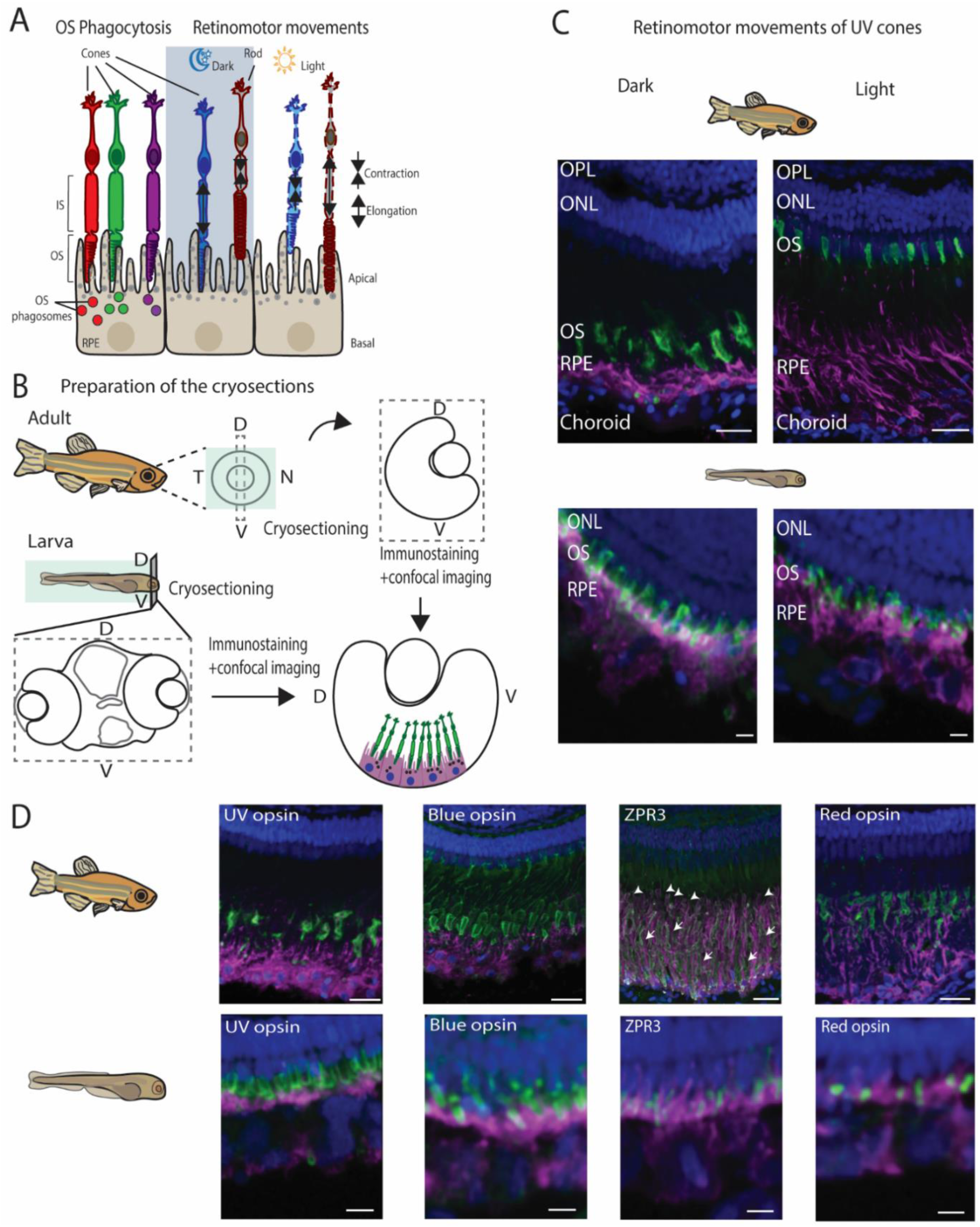
Adult and larval zebrafish retinal cryosections show structural differences at the photoreceptor-RPE interface. A) Illustrative figure of the RPE-photoreceptor interface, outer segment (OS) phagocytosis and OS movement during the retinomotor movements in dark and light. Not to scale. B) Illustrative figure of preparation and immunolabelling of adult and larval transverse retinal cryosections along the dorso-ventral axis. Not to scale. D: dorsal, V: ventral, N: nasal; T: temporal. C) Confocal images of adult (up) and larval (below) zebrafish retinal cryosections showing the interface between the RPE and UV cones in dark and light conditions. Adult cryosections show robust retinomotor movements seen as elongation and contraction of the UV cone OSs in the dark and light, respectively. Similar movements are not seen in larval cryosections (scale bars in the adult sections 20 µm; in the larval sections 5 µm). D) Confocal images of adult (upper row) and larval (lower row) showing immunolabelling of each cone subtypes together with the RPE. Scale bars in the adult sections 20 µm and in the larval sections 5 µm. Green: outer segments of the photoreceptors, Magenta: RPE, Blue: nuclei. IS: photoreceptor inner segment, ONL: outer nuclear layer, OPL: outer plexiform layer, OS: outer segment, RPE: retinal pigment epithelium

In the present study, we investigated the rhythmicity of the photoreceptor OS phagocytosis of different cone subtypes using young cone-dominant zebrafish larva as an animal model. Our results demonstrate ongoing phagocytosis of all cone subtypes with differences in the rhythmicity, which are likely due to the asymmetry in their maturation states. We also examined the effect of the ambient light on the process in these animals. Our results show that removal of the external light suppresses the rhythmic nature of the OS phagocytosis indicating that the daily peaks of phagocytosis are primarily controlled by the ambient light in developing zebrafish.

## 2. Materials and Methods

### 2.1 Zebrafish husbandry and maintenance

Wild type zebrafish (*Danio rerio*) larvae at 7 *dpf* were used in the experiments as they have fully functional cone cell-mediated perception. The embryos and larvae were grown in 10 cm petri dishes in E3 embryo solution (5 mM NaCl, 0.17 mM KCl, 0.33 mM CaCl_2_, 0.33 mM MgSO_4_, 0.0003 g/l Methylene Blue, pH 7.2) supplemented with 0.0045 % 1-Phenyl 2-thiourea (PTU, Sigma-Alrich, St. Louis, MO, USA) and maintained at 28.5 °C in an incubator. Larvae were raised under normal 14 h/10 h light-dark cycle (LD), unless otherwise stated and were fed once a day with GEMMA Micro 75 (Skretting, Stavanger, Norway). Male and female animals were randomly chosen to each experimental group. For sample collection, zebrafish larvae were euthanized with an overdose (0.08 %) of an anesthetic Tricaine (3-aminobenzoic acid ethyl ester, pH 7.0) (Sigma-Aldrich, USA) prepared in E3 embryonal solution.

1- to 1.5-years-old adult Albino (slc45a2^ti225^) zebrafish (European Zebrafish Resource Center, Eggenstein-Leopoldshafen, Germany) were used for making histological cryosections. The fish were housed in flow-through water-circulation system (at 25 °C) under LD. Male and female animals were randomly chosen for cryoblock preparation. Fish were euthanized with an overdose (0.08 %) of Tricaine.

### 2.2 Ethics statement

The wild type zebrafish (*Danio rerio*) with AB/Tübingen background line, as well as the homozygote Albino (*slc45a2^ti225^*) with Tübingen background line were obtained from Tampere Zebrafish Core Facility (Tampere University, Finland). Zebrafish were maintained according to the standard protocols and the ethical guidelines set by the Ethical Board in Finland. The husbandry and the experiments were done in accordance with the Finnish Act on the Protection of Animals Used for Scientific or Educational Purposes (497/2013) and the EU Directive 2010/63/EU. The ethical permissions for the experiments were granted by the Animal Experiment Board in Finland (the licenses ESAVI/37052/2020 and ESAVI/46385/2023).

### 2.4 Light cycle experiments and sample collection

To study the rhythmicity of OS phagocytosis, 7 *dpf* old larvae that were raised in 28.5 °C in an incubator with normal light cycle (14h light/12 h dark; LD), were used. During the time point experiments, at 6 *dpf,* the larvae were transferred to room temperature (RT) at least 24 h prior to sample collection to match the environmental temperature with the temperature at which later dark-adaptation experiments were performed. To maintain LD light cycle at RT, an extra spotlight was utilized in the lab, and for the dark period, the larvae were placed in an opaque box. Larval samples were collected and euthanized at seven different time points over a 24 h period following light onset. Sample collection time points are expressed in Zeitgeber time (ZT), and were as follows: ZT1, ZT3, ZT5, ZT10, ZT16, ZT18, ZT23 (Figure 3B), where ZT0 is the light onset and ZT14 light offset. Larvae that were collected during the dark period, at ZT16, ZT18 and ZT23, were euthanized in a dark room to avoid influence of external light. Furthermore, to study the effect of external light on the rhythmicity of OS phagocytosis, a separate group of 6 *dpf* larvae were dark adapted at least 24 h prior to sample collection. These larvae were kept in dark-dark cycle (DD) by placing them into an opaque box in a dark room at RT. Larvae were collected at the same seven time points (ZT1, ZT3, ZT5, ZT10, ZT16, ZT18, ZT23) as previously described and euthanized in dark room during sample collection.

### 2.5 Cryoblock preparation and cryosectioning

For cryoblock preparation of adult zebrafish samples, the fish were euthanized with Tricaine prior to enucleation. Collected eyes were then immersion-fixed in 4% paraformaldehyde (Sigma-Aldrich, USA) in phosphate-buffered saline (PBS) for 2 h at RT followed by three PBS washes of 5 minutes (min) each. For cryoblock preparation from larval zebrafish, larvae were collected at the time points mentioned above and euthanized with Tricaine prior to immersion-fixation with 4 % paraformaldehyde in PBS for 2 h at RT or overnight at +4 °C followed by three 5 min PBS washes. After fixation, larvae and adult zebrafish eyes were sucrose-protected by incubating the samples in sucrose gradient with rising concentrations (5 %, 10 %, 20 %, 30 %) for 1 h in each solution at RT, except 30 % sucrose solution in which the incubation lasted for 24 h at +4 °C. After sucrose-protection, the larvae and adult eyes were embedded in tissue-freezing medium (Tissue-Tek O.C.T compound, Sakura Finetek, Torrance, CA, USA) and frozen with liquid nitrogen. Larval cryoblocks were then sectioned with a cryotome (MEV+ cryostat, SLEE medical GmbH, Nieder-Olm, Germany) along dorso-ventral axis of the larval head to obtain 10 µm thick cross-sections including both larval eyes. Similarly, enucleated adult eye cryoblocks were sectioned to obtain 10 µm thick eye cross-sections. Cryosections were attached to adhesive microscope slides (Epredia Superfrost Plus Adhesion Microscope slides, Fischer Scientific, Waltham, MA, USA) and incubated at 60-62 °C for 2 h to ensure proper drying and attachment to the slides.

### 2.6 Immunohistochemistry

Cryosections were immunolabelled for opsin type- and RPE tissue-specific proteins to observe phagocytosed OS phagosomes from certain cone type inside the RPE tissue. The binding sites of the used antibodies are listed in the supplementary table (Supplementary S1). Prior to immunolabelling, the sections were washed with 1x Tris-Buffered Saline (TBS)-Tween (0.05 % of Tween) and permeabilized by incubating them in 0.05 % Triton-x-100 in TBS-Tween for 15 min and subsequently blocked with 3 % Bovine Serum Albumin (BSA) (Sigma-Aldrich) in TBS for 1 h at RT. All primary and secondary antibody dilutions were prepared in 3 % BSA-TBS. Primary antibodies with the following concentrations were used in this study: anti-zebrafish Blue opsin (1:300), anti-zebrafish UV-opsin (1:300), anti-rhodopsin [1D4] (1:100), zpr-3 (1:200), Anti-RPE65 [N1C3] (1:300), zpr-2 (1:200), anti-rod opsin (1:200) (Table 1). Cryosections were incubated with primary antibodies for 24 h at +4 °C followed by two TBS-Tween washes and 24 h incubation with secondary antibodies (1:200) and phalloidin (1:100) (Table 2) in the dark. After secondary antibody incubation, the samples were washed twice with TBS-Tween and once with MilliQ water followed by DAPI in MilliQ water (1:1200) labelling for 8 min at RT in the dark. Prior to mounting the samples with ProLong Diamond antifade mounting medium (Thermo Fischer, Waltham, MA, USA), the samples were washed once with MilliQ water.

**Table 1.** Primary antibodies with their target cell types, manufacturers, Research Resource Identification (RRID) numbers and animal species in which the antibodies are produced.

| Primary antibodies |  |  |  |  |  |
| --- | --- | --- | --- | --- | --- |
| Antibody | Labelled cells | Manufacturer | Catalog number | Animal | RRID numbers |
| UV opsin | UV cones | Kerafast<br>(Boston, MA, USA) | EJH013 | rabbit |  |
| Blue opsin | Blue cones | Kerafast<br>(Boston, MA, USA) | EJH012 | rabbit |  |
| Rod opsin | Rods | Sigma-Aldrich<br>(St. Louis, MO, USA) | O4886 | mouse | AB_260838 |
| Rhodopsin [1D4] | Red cones in zebrafish<br>(Yin et al., 2012) | Abcam<br>(Cambridge, UK) | ab5417 | mouse | AB_304874 |
| zpr-3 | Green cones+Rods<br>(Hu et al., 2024) | ZIRC<br>(Eugene, OR, USA) |  | mouse | AB_10013805 |
| RPE65 | RPE | GeneTex | GTX103472<br>(Irvine, CA, USA) | rabbit | AB_2037911 |
| zpr-2 | RPE | ZIRC<br>(Eugene, OR, USA) |  | mouse | AB_10013804 |

**Table 2.** Secondary antibodies, phalloidin and DAPI with their targets, manufacturers and Research Resource Identification (RRID) numbers.

| **Secondary antibodies, phalloidin and DAPI** |  |  |  |  |
| --- | --- | --- | --- | --- |
| **Antibody** | **Target** | **Manufacturer** | **Catalog number** | **RRID number** |
| Goat anti-Mouse IgG (H+L) Highly Cross-Adsorbed Secondary Antibody, Alexa Fluor™ 488 | mouse primary antibodies | Thermo Fischer Scientific (Waltham, MA, USA) | A-11029 | AB_2534088 |
| Goat anti-Rabbit IgG (H+L) Cross-Adsorbed Secondary Antibody, Alexa Fluor™ 488 | rabbit primary antibodies | Thermo Fischer Scientific (Waltham, MA, USA) | A-11008 | AB_143165 |
| Goat anti-Rabbit IgG (H+L) Cross-Adsorbed Secondary Antibody, Alexa Fluor™ 568 | rabbit primary antibodies | Thermo Fischer Scientific (Waltham, MA, USA) | A-11011 | AB_143157 |
| Goat anti-Mouse IgG (H+L) Highly Cross-Adsorbed Secondary Antibody, Alexa Fluor™ 568 | mouse primary antibodies | Thermo Fischer Scientific (Waltham, MA, USA) | A-11031 | AB_144696 |
| Phalloidin 643 | F-actin (cell cytoskeleton) | ATTO-TEC (Siegen, Germany) | AD643-81 |  |
| 4',6'-diamidine-2-phenylindole dihydrochloride (DAPI) | DNA (nuclei) | Sigma-Alrich, St. Louis, MO, USA | 10236276001 |  |

### 2.7 Confocal microscopy and image processing

All samples were imaged with Nikon A1R laser scanning confocal microscope mounted in inverted Nikon Ti-E body (Nikon, Tokyo, Japan). Nikon Apo 40x/1.15 DIC (Water) objective was used to image larval whole eye cryosections and Nikon Apo 60x/1.40 DIC (Oil) objective was used to image specific areas of larval and adult eye cryosections. 1024 × 1024 pixel large Z-stacks were taken to image the whole eyes (larvae) or specific retinal areas (adults and larvae) using 405 nanometer (nm), 488 nm, 561 nm, 640 nm channels for fluorescent DAPI, RPE-specific protein, photoreceptor-specific opsin proteins and phalloidin, respectively.

The imaging data was saved in .czi format and images were processed with ImageJ-Fiji-64 bit software. Only linear brightness and contrast adjustments were performed for the pixel intensities in the images. Gaussian filtering with radius 1 (σ= 1) was done to improve the image quality when required. Final figures and graphs were constructed with Adobe Illustrator.

### 2.8 Analysis tool for quantitative analysis of OS phagosomes

The detection of OS phagosomes was performed using a custom-developed semi-automatized analysis tool for ImageJ-Fiji (ImageJ 1.54) (Supplementary S2). The tool utilizes standard ImageJ-Fiji libraries for general image processing and MorphoLibJ (MorphoLibJ_-1.6.2) for all morphological operations.

The analysis tool takes confocal z-stack images of fluorescently labeled cryosections containing RPE and photoreceptor channels as input. Before further analysis, the stacks are processed into maximum intensity projections and denoised using a Gaussian filter (σ= 1). The entire workflow was executed as an interactive script that allows the user to adjust parameters as needed. First, a user-defined RPE area is binarized through auto-thresholding using Li’s method (Li and Lee 1993). A morphological opening with a disc-shaped structuring element of radius 6 pixels is then applied to remove holes and smooth the selection. The final selection is saved as an ImageJ region of interest (ROI) to be used as the area to search for OS phagosomes in subsequent steps. OS phagosome segmentation is based on the extraction of local bright objects using the h-dome transform, first introduced by Vincent (1993) and since used for spot-detection tasks in fluorescent and electron microscopy (EM) datasets (Smal et al. 2010; Geng et al. 2023), to detect all local maxima. h-dome transform method suppresses low-contrast regions and enhances significant intensity peaks bigger than defined h-value. The Maxima Finder is then applied to detect the intensity peaks with an additional criterion called prominence that is the minimum height difference between a local maximum and its surrounding region for the maximum to be considered significant.

For analysis tool development and parameter testing, a test set of Z-stacks with each cone subtype labelled individually along with the RPE tissue was used. Empirical testing was conducted to determine appropriate values for the parameters h-value and prominence to find the balance between sensitivity (detecting enough peaks) and specificity (avoiding false positives or irrelevant peaks) (Supplementary S2). The test analysis for each image was performed with the analysis tool using all combinations of five h-dome values (5, 15, 25, 50, 100) and five prominence thresholds (5, 10, 20, 40, 80). The local maxima detected by the analysis tool were then matched to the manually annotated local maxima (Ground truth) and the detected peaks were defined as True positive (TP), False positive (FP) and False negative (FN) as follows: A detected peak was considered a TP if it was within three pixels of a ground truth peak, a detected peak that could not be mapped to any ground truth peak was classified as a False Positive (FP), and a ground truth peak that was not mapped to any detected peak was classified as a False Negative (FN). By using the counts of TP, FP and FN, the values for Precision, Sensitivity, F1-score and False positive rate (FPR) (Defined in Supplementary S2) were calculated for each parameter combination of h-values and prominence thresholds to evaluate the algorithm’s performance and its overall detection reliability in the current analysis (Supplementary S2).

Since the parameter combinations of all five h-dome values and prominence thresholds between 5-20 produced relatively similar values for precision, sensitivity, F1-score and FNR (Supplementary S2), the performance and peak detection reliability with these parameters were highly equal. Therefore, in our analysis tool, the h-value was set to 15 and the prominence threshold was set specifically (always under the value of 20) for each analysed image during the OS phagosome analysis to reach the most reliable detection of OS phagosomes. All analysis tool results were carefully inspected, and when the tool failed to detect the appropriate number of OS phagosomes, the number of phagosomes was quantified manually.

### 2.9 Quantitative analysis and statistical testing

For OS phagosome number analysis, 4-12 larval whole eye cryosections (indicated as “n” in the figure legends) from total of 3-9 larval individuals per time point were analysed so that each section represented individual larval eye. The OS phagosomes found in the RPE-tissue were quantified from the whole eye cryosections using the custom-developed analysis tool described above. The numbers of OS phagosomes found in the RPE tissue were normalized to the RPE length, which was measured from the RPE-OS interface individually in each cryosection (Figure 2B). The data values in the graphs 3C and 4B show the number of OS phagosomes per 10 µm of RPE as well as the mean for the total number of samples (Figure 3C, Figure 4B). Error bars represent standard errors of the mean (SEM). Normality of data distributions were tested using the Shapiro-Wilk test.

**Figure 2.**
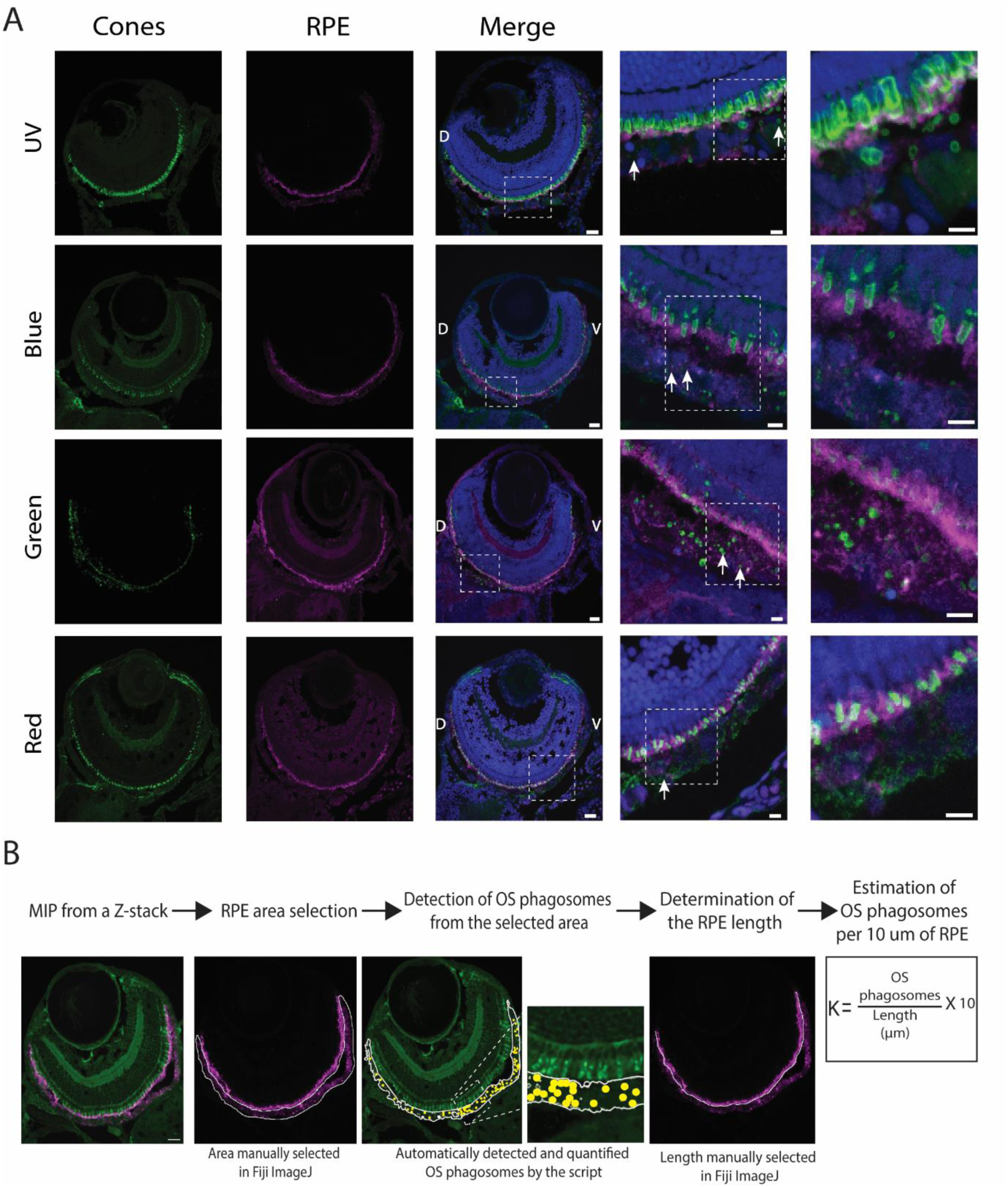
OS phagosomes from all cone subtypes can be detected and quantified from cryosections of the 7 dpf old zebrafish larvae. A) Immunolabelling of cone subtype-specific opsins and RPE tissue in the 7 dpf larval cryosections reveal OS phagosomes (white arrows) inside the RPE verify that OSs of all subtypes (UV, blue, green and red) are phagocytosed in the developing zebrafish. Scale bars in the whole eye cryosections 20 µm and 5 µm in the zoomed-in views marked as white dashed boxes. B) Operational steps of the semi-automatized analysis tool for quantification of OS phagosomes from the selected RPE area in the larval cryosections as well as defining the number of OS phagosomes per 10 µm of RPE tissue. Yellow circles represent the detected OS phagosomes. RPE: retinal pigment epithelium, D: dorsal, V: ventral, MIP: maximum intensity projection, OS: outer segment, K: Number of OS phagosomes per 10 µm of RPE.

**Figure 3:**
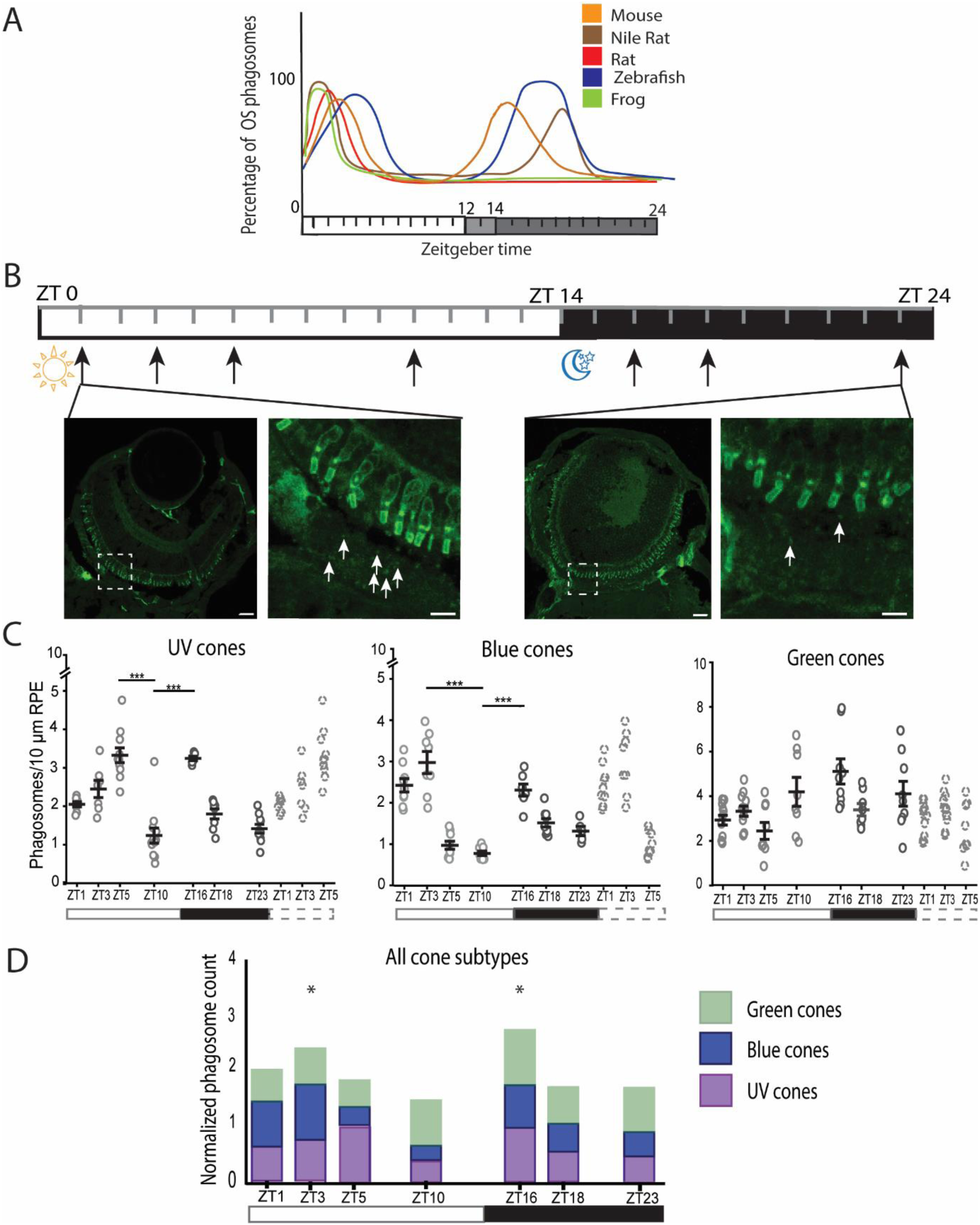
Phagocytosis of the UV and blue cone OSs shows two phagocytic peaks during the 24 h cycle in the zebrafish larvae. A) Illustrative figure of previous data generally demonstrates one or two activity peaks of OS phagocytosis for rod-dominant and cone-dominant species, respectively. Figure is created after Moran et al., 2022. B) 7 dpf zebrafish larvae were collected at seven time points (black arrows) and used for the cryosection preparation. After immunolabelling the sections for cone-subtype specific opsins, the numbers of OS phagosomes were quantified. The differences in the numbers of phagosomes from blue cone OSs (white arrows) at two time points, ZT1 and ZT23, were evident in the zoomed-in views (areas marked with dashed boxes). Scale bars in the whole eye sections are 20 µm and in zoomed-in views 5 µm. C) Scatter plots show quantitative results of phagosome numbers of UV, blue and green cone OSs during the 24 h cycle. White and black bars show the light (ZT0-ZT14) and dark (ZT14-ZT24) phases of the day, respectively. Dashed bar show the repeated three first sample collection time points. Phagocytosis of UV and blue cone OSs showed rhythmic nature and peaked two times per 24 h, whereas green cone OSs were phagocytosed more constantly throughout the day. Data shows the mean ± SEM. *p < .05, **p < .01, ***p < .001. D) Stacked bar graph containing normalized data from UV, blue and green cones show two phagocytic peaks (marked with asterisks), and that UV cones and blue cones affect more the rhythmic profile of phagocytosis activity than green cones. ZT: zeitgeber time, OS: outer segment, LD: Light-dark condition.

**Figure 4:**
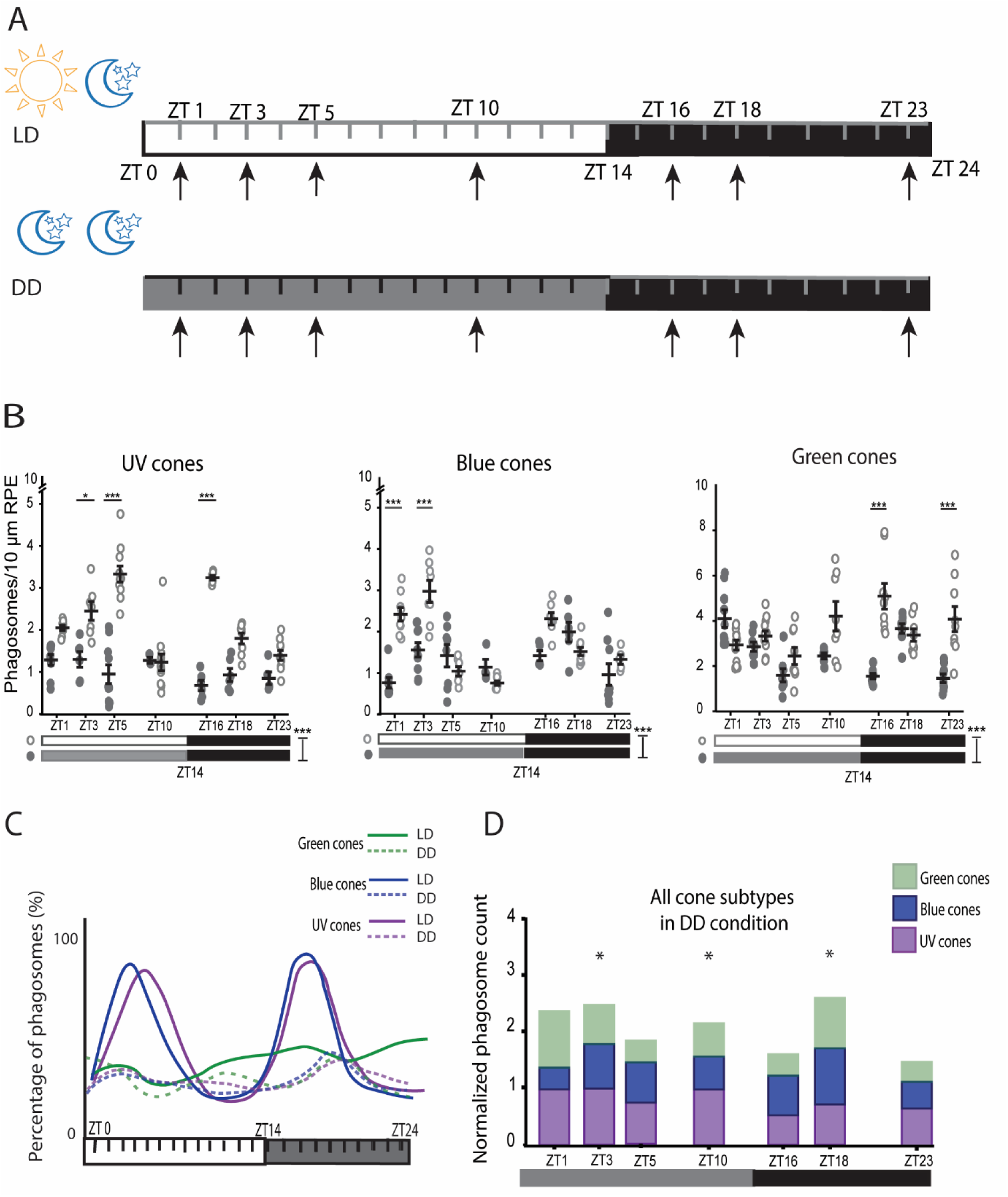
The peaks of UV and blue cone OS phagocytosis dampen in constant darkness. A) Zeitgeber timelines showing sample collection times (black arrows) in normal light cycle, LD (upper) and in constant darkness, DD (lower). B) Scatter plots show phagosome numbers from UV, blue and green cone OSs during the 24 h cycle in LD (light circles) and in DD (dark circles) cycles. In addition, the peaks in the numbers of phagosomes from UV and blue cone OSs significantly dampened in DD compared to LD. Data shows the mean ± SEM. *p < .05, **p < .01, ***p < .001. C) Illustrative figure of the peaks of UV, blue and green cone OS phagocytosis in LD and DD shown as percentages of OS phagosomes found in the RPE over the 24 h cycle. The clear peaks of UV and blue cone OS phagosomes diminished in DD condition, and the number of OS phagosomes from all three cone subtypes decreased in DD compared to LD condition. D) Stacked bar graph containing the normalized data from UV, blue and green cones shows slight rhythmicity in constant darkness with three time points at which the number of phagosomes increased (asterisks). ZT: Zeitgeber time, OS: outer segment, LD: Light-dark condition, DD: Dark-dark condition

#### LD cycle experiments

Data for each cone subtype was considered as normally distributed. One-way ANOVA was performed to determine whether there was a statistically significant difference in the mean of OS phagosome numbers across the time points (ZT1, ZT3, ZT5, ZT10, ZT16, ZT18, ZT23) (Figure 3). Subsequent Bonferroni Post Hoc test was performed to determine which of the time points differed from each other. p value <0.05 was considered as statistically significant. Finally, the OS phagosome numbers at the peak time points were compared to the baseline OS phagosome number to evaluate whether the differences were statistically significant (Figure 3C). For this study, the baseline time point (ZT10) was chosen based on previous data by Lewis et al. (2018) that showed lowest number of phagosomes at ZT10.5.

#### LD vs DD experiments

Data in LD and DD was considered as normally distributed for each cone subtype. A two-way ANOVA was performed to examine the effect of light condition (LD versus DD) over the time points (ZT1, ZT3, ZT5, ZT10, ZT16, ZT18, ZT23) on the numbers of OS phagosomes found in the RPE (Figure 4B). Subsequent Bonferroni Post Hoc test was performed to determine which of the time points differed from each other in their OS phagosome numbers between LD and DD conditions. Finally, figures 3D and 4C show the bar graphs of combined phagosome numbers for UV, blue and green cone OSs in LD and DD, respectively. For these, the numbers of OS phagosomes from each cone subtype were normalized to the level of the highest number of phagosomes (phagosomes/10 µm of RPE) over the studied time points. Then, the normalized values for UV, blue and green cones were summed together. All statistical tests in this study were performed with Origin 2019b 64 bit and IBM SPSS Statistics 29.0.1.0. Final graphs were created with Origin and Adobe Illustrator.

## 3. Results

### 3.1. Adult and larval zebrafish retinal cryosections show differences in the photoreceptor-RPE interactions

The organization of photoreceptors is well characterized in both adult and larval zebrafish retinae. In adults, OSs of different photoreceptor types are organized into different layers, whereas in larvae, similar multilayer organization of photoreceptors is absent (Allison et al., 2010; Branchek & Bremiller, 1984, Supplementary S3). Additionally, in adults, the interaction between photoreceptors and the RPE changes significantly in different light conditions and during circadian-related processes, such as OS tip phagocytosis and retinomotor movements containing elongation and contraction of rods and cones (Lakkaraju et al., 2020; Menger et al., 2005, Figure 1A). Instead, in larvae under 28 *dpf*, retinomotor movements of photoreceptors are minimal (Hodel et al. 2006) and their phagocytosis has not been thoroughly studied.

To investigate and compare OS phagocytosis in adults and larvae, we prepared histological cross-sections along dorso-ventral axis from both adult and larval zebrafish eye cryoblocks (Figure 1B). Next, we immunolabelled RPE and photoreceptor cells in the sections by RPE-specific antibody and photoreceptor-type specific opsin-antibodies (Figure 1B). We used 7 *dpf* old larvae for our studies, as all cone subtypes (UV, blue, green and red) are detectable and functional already at 5 *dpf* (Branchek and Bremiller 1984; Saszik et al. 1999; Crespo and Knust 2018). In accordance with previous studies (Menger et al. 2005; Hodel et al. 2006), our confocal images show robust and clearly observable retinomotor movements of photoreceptors in adult, but not in the 7 *dpf* old larval zebrafish retina (Figure 1C). Indeed, the adult UV cone OSs extended more into the RPE tissue in the dark than in the light (Figure 1C, upper row), but similar movements were not seen in the larvae (Figure 1C, lower row). With respect to OS phagocytosis studies, multilayered organization and retinomotor movements make it challenging to distinguish the OS phagosomes from the intact photoreceptor outer segment layer in the adult fish. Moreover, estimating the exact localization of the OS phagosomes inside the RPE at different time points is demanding as it is hard to determine whether they are in the long apical protrusion structures or inside the cell body. Since the retinomotor movements are lacking from larval zebrafish, for our approach with cryosections and immunohistochemical labelling, larval zebrafish is a more suitable model.

In search of specific antibodies to the cone subtypes in zebrafish, we found no publicly available specific antibodies for green cone opsin, in contrast to antibodies specific for the other opsins. Therefore, to detect green cones, we used zpr-3, an antibody that is shown to label OSs of green cones and rods in adult zebrafish (Hu et al. 2024). Our confocal images of adult and larval zebrafish cryosections demonstrate that antibodies for UV opsin, blue opsin and red opsin label only one retinal layer as they specifically target UV, blue and red cones, respectively (Figure 1D, Supplementary 3). On the contrary, adult zebrafish cryosections clearly display zpr-3 labelling in two retinal layers demonstrating rod and cone OSs (Figure 1D). Rod OSs (arrows) highly overlap with the RPE microvilli that extend up to 60 µm from the nuclear layer of the RPE, whereas the green cone OSs (arrowheads) locate closer to the inner retina. However, cryosections of 7 *dpf* larval zebrafish exhibit zpr-3 labelling only in one retinal layer (Figure 1D, Supplementary 3). Since at this age fully functional rods are not formed yet (Branchek and Bremiller 1984), we believe the antibody reveals principally only OSs of the green cones.

The challenges faced with the rod/green cone labelling by zpr-3 antibody, as well as with the OS phagosome detection caused by retinomotor movements in adult zebrafish can be overcome using young zebrafish larvae. This makes larval zebrafish a potential model for our studies on phagocytosis of individual cone subtype OSs. In addition, since the phagocytosis process is necessary for the photoreceptor viability and for normal visual function, we investigated whether the rhythmicity of this process is developed already in young zebrafish larvae, where the organization of the photoreceptors and the entire retina are still under modification.

### 3.2 All Cone subtypes are phagocytosed in the 7 *dpf* old zebrafish larva

A previous study by Lewis et al. (2018) showed evidence of cone phagocytosis in 7 *dpf* old zebrafish larvae, but phagocytosis of the different cone subtypes has remained unexplored. Here, to investigate whether all the different cone subtypes are being phagocytosed in young zebrafish larvae, we used the above-described approach with cryosections, immunohistochemical labelling and imaging to detect OS phagosomes, originating from different cone subtypes, in the RPE. Our confocal microscopy data shows that phagosomes from UV, blue and green cone OSs can be detected in the RPE (Figure 2A). Moreover, the phagosomes from the red cone OSs were observed in the RPE, although the red opsin label in the images had more background signal than the other opsin labels, making it more challenging to distinguish the OS phagosomes from the background. Nevertheless, our data verifies that all cone subtypes are phagocytosed already in 7 *dpf* old larvae (Figure 2A).

To evaluate the level of phagocytosis at different time points, the cone phagosomes were quantified from the cryosections. To facilitate efficient quantification of the OS phagosomes, we developed a semi-automatized analysis tool (Supplementary 2). Engulfed OS particles are highly heterogeneous in size, shape, and fluorescence intensity, due to their state of degradation by the RPE’s endo-lysosomal machinery. This heterogeneity posed a challenge for automated detection. Moreover, the unspecific labeling of surrounding tissues lowered the contrast between particles and the background, making manual quantification highly demanding. We selected a semi-automatized approach to avoid the need for tedious training of more intricate detection systems and to allow easy adaptation of the workflow to different cone subtype samples. For that, we developed an analysis tool that uses confocal z-stacks of fluorescently labelled larval whole-eye cryosections containing RPE and photoreceptor channels as input. For an OS particle to be considered as phagocytosed, its label had to overlap with the RPE label. This requirement effectively divided the rest of the workflow into two sections: RPE area selection and OS phagosome segmentation (Figure 2B, supplementary 2). After segmenting the RPE area manually, the analysis tool detected and quantified OS phagosomes found in that area. Figure 2B also shows how the total length of the RPE tissue in the images was manually defined and how the number of OS phagosomes (K) per 10 µm of RPE tissue was calculated in later experiments.

To verify the accuracy and specificity of the analysis method to detect the OS phagosomes within the selected RPE area, the OS phagosome numbers were counted also manually from the images. The numbers of OS phagosomes for UV, blue and green cones were comparable when quantified by either way (maximum difference in the phagosome numbers 10.5 %) (Table 3). Thus, the analysis tool showed its reliability in quantification of OS phagosomes from immunofluorescent labelled cryosections. However, the tool did not work optimally for the red opsin label, since the numbers of OS phagosomes differed notably when quantified manually (minimum difference in the phagosome numbers 58 %, maximum difference 129 %) (Table 3). Therefore, we did not use the analysis tool further for the quantification of red cone phagosomes, but instead, later quantifications were performed manually.

**Table 3.**
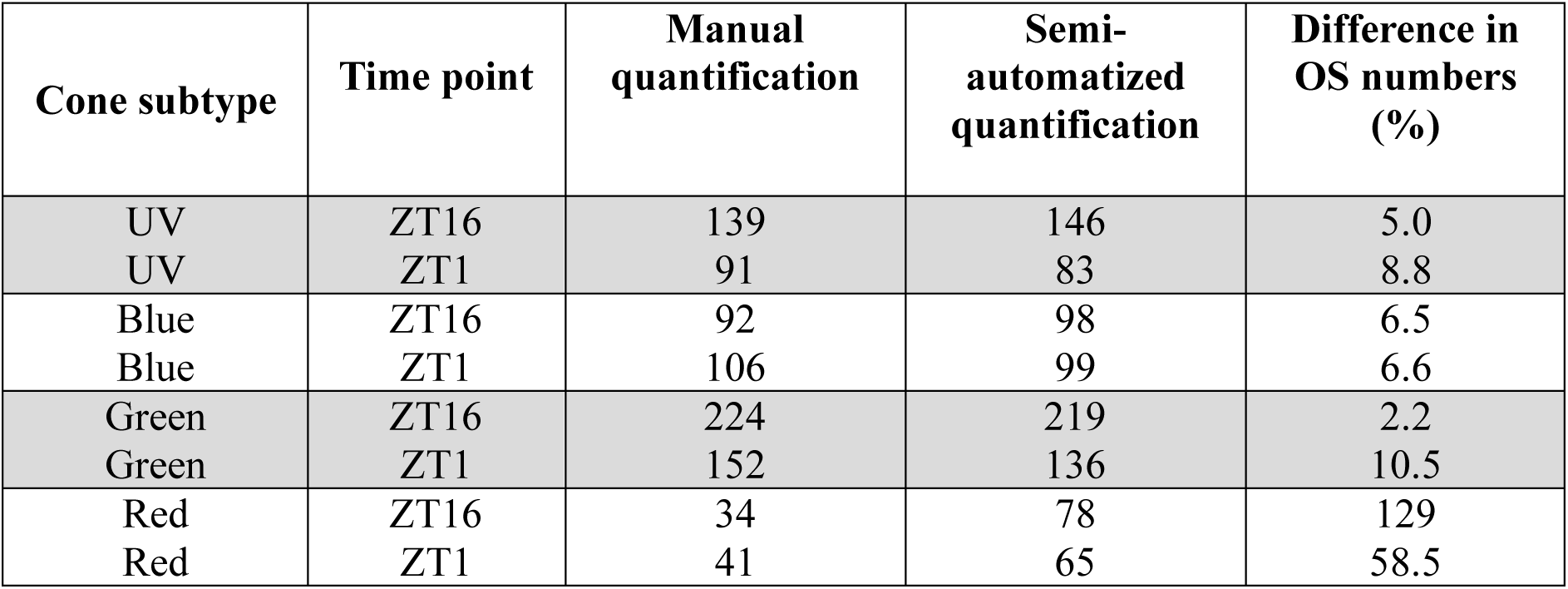
Comparison of the total OS phagosome numbers from different cone subtypes at two time points quantified manually and using the semi-automatized analysis tool from the larval zebrafish cryosections. OS: outer segment, ZT: Zeitgeber time.

| Cone subtype | Time point | Manual quantification | Semi-automatized quantification | Difference in OS numbers (%) |
| --- | --- | --- | --- | --- |
| UV | ZT16 | 139 | 146 | 5.0 |
| UV | ZT1 | 91 | 83 | 8.8 |
| Blue | ZT16 | 92 | 98 | 6.5 |
| Blue | ZT1 | 106 | 99 | 6.6 |
| Green | ZT16 | 224 | 219 | 2.2 |
| Green | ZT1 | 152 | 136 | 10.5 |
| Red | ZT16 | 34 | 78 | 129 |
| Red | ZT1 | 41 | 65 | 58.5 |

### 3.3 Phagocytosis of UV and blue cones show rhythmicity in 7 *dpf* old zebrafish larvae

The activity of OS phagocytosis peaks once or twice per day depending on the animal species (Reviewed by Moran et al. 2022b, Figure 3A). To study the rhythmicity of OS phagocytosis of the different cone subtypes under normal LD conditions, zebrafish larvae were collected at seven discrete time points shown at ZT timeline with black arrows (Figure 3B). The samples were prepared and imaged as previously explained. Figure 3B also shows larval whole-eye section immunolabelled with blue cone subtype-specific antibody at two different time points ZT1 and ZT23. The difference in the OS phagosome numbers is clearly visible in the images between the time points with more OS phagosomes at ZT1 compared to ZT23 (Figure 3B).

Phagosomes from UV, blue and green cone OSs were quantified from the defined RPE area in the images using the developed analysis tool. As the tool did not work reliably with the red cones, the phagosomes from red cone OSs were quantified manually. Since the detection of these phagosomes from the background labelling was challenging, we consider red cone data more unreliable than the data of the other cone subtypes. Therefore, we concentrated on UV, blue, and green cones in our further study, and red cone data is shown only as supplementary information (Supplementary S4, Supplementary S6). Our data shows that the numbers of OS phagosomes varied significantly between the distinct cone subtypes as well as between the discrete time points (Figure 3C, supplementary S4). We used one-way ANOVA analysis to show statistically significant differences in phagosome numbers from UV, blue and green cones over the 24 h (Table 4) (p= 1.69*10^-14; p=9.10*10^-15; p=7.75*10^-4, respectively). Subsequent Bonferroni Post Hoc test revealed two time points for UV (ZT5 and ZT16) and blue cones (ZT3 and ZT16) at which the OS phagosome numbers were significantly increased (Figure 3C). The number of phagosomes from UV cone OSs gradually increased first after light onset peaking at ZT5 (3.33 phagosomes/10 µm of RPE), and secondly after light offset at ZT16 (3.25 phagosomes/10 µm of RPE). At both peaks, the number of phagosomes from UV cone OSs was significantly higher compared to the baseline at ZT10 (baseline 1.24 phagosomes/10 µm of RPE, differences p= 1.22*10^-12 for ZT5 and p= 1.74*10^-9 for ZT16). Similarly, the second peak of phagosomes from blue cone OSs appeared after light offset at ZT16 (2.29 phagosomes/10 µm of RPE), but the light onset-associated peak occurred earlier, at ZT3 (2.97 phagosomes/10 µm of RPE). At both peaks, the numbers of OS phagosomes were significantly higher compared to the baseline at ZT10 (baseline 0.77 phagosomes/10 µm of RPE, differences p=1.44 *10^-12 for ZT3 and p=1.94*10^-7 for ZT16). Instead, the numbers of phagosomes from green cone OSs showed less variation between the time points, and clear peaks were not observed (Figure 3C). This indicates that green cones are phagocytosed at a more constant level throughout the 24 h cycle compared to the other subtypes. Moreover, the green cone data showed higher numbers of OS phagosomes at each time point compared to the other subtypes (Figure 3C, Table 4). Interestingly, manually calculated phagosomes from red cone OSs showed only one peak that emerged at a late dark timepoint of the day, at ZT23 (1.93 phagosomes/10 µm of RPE), with significantly more OS phagosomes compared to the baseline at ZT10 (baseline 0.69 phagosomes/10 µm of RPE, difference p= 1.80*10^-6) (Supplementary S4). In addition, the total number of phagosomes from red cone OSs throughout the 24 h cycle was lower than that of the other cone subtypes (Table 4, Supplementary S4). Lastly, in addition to cone immunolabelling, a set of larval cryosections were immunolabelled with anti-rod opsin antibody, previously shown to label rods in various vertebrate species such as mouse, rat, duck, turtle and goldfish (Barnstable 1980, Silver et al. 1988) to visualize rod precursors and evaluate the level of their phagocytosis by the RPE. Surprisingly, our results show that the rhythmicity profile of rod phagocytosis corresponds well to that of green cones (Supplementary S5).

**Table 4.** Averages of phagosomes from UV, blue and green cone OSs per 10 µm of RPE at each time point. n≥5 sections at each time point, each section represents one larva. OS: outer segment.

| **Time** | **UV cones** | **Blue cones** | **Green cones** | **Average from the sum of UV, blue and green cone OSs** |
| --- | --- | --- | --- | --- |
| ZT1 | 2.05 | 2.42 | 2.93 | 2.53 |
| ZT3 | 2.45 | 2.97 | 3.33 | 3.02 |
| ZT5 | 3.33 | 0.97 | 2.45 | 2.38 |
| ZT10 | 1.24 | 0.77 | 4.18 | 1.88 |
| ZT16 | 3.25 | 2.29 | 5.10 | 3.65 |
| ZT18 | 1.80 | 1.50 | 3.38 | 2.22 |
| ZT23 | 1.41 | 1.31 | 4.09 | 2.36 |
| Average of total OS phagosome count over 24 hours | 2.15 | 1.76 | 3.59 |  |

To evaluate the phagocytosis rhythmicity profile of each cone subtype in relation to the combined cone OS phagocytosis profile, automatically quantified phagosome numbers (i.e. from UV, blue and green cone OSs) were normalized to the level of the highest number of OS phagosomes (phagosomes/10 µm of RPE) over the studied time points individually for each cone subtype. The stacked bar graph containing normalized values shows that the sum of the phagosomes from UV, blue and green cone OSs peaked twice during the 24 h cycle (Figure 3D), which is in line with the previous zebrafish studies (Lewis et al. 2018; Moran et al. 2022a). The first peak was associated with light onset and occurred at ZT3, whereas the second peak occurred after light offset at ZT16. The data also indicates larger impact of UV and blue cones than green cones on the overall phagocytosis rhythmicity profile as the UV and blue cone bars as well as their combination form one light-associated and one dark-associated peak (Figure 3D). The non-normalized values of phagosomes from UV, blue and green cone OSs were then summed together at each time point and after averaging, the numbers of phagosomes at the peak time points were 3.02 phagosomes/10 µm of RPE and 3.65 phagosomes/10 µm of RPE, at ZT3 and ZT16, respectively (Table 4). Table 4 shows that at both peak time points, the number of phagosomes was higher compared to the baseline at ZT10 (1.88 phagosomes/10 µm of RPE). Collectively, our results indicate that the phagocytosis of larval UV and blue cone OSs, but not green OSs, is rhythmically regulated.

### 3.4 The rhythmic nature of the UV and blue cone phagocytosis reduces in constant darkness

Previous studies with constant dark environment have revealed that rhythmicity of OS phagocytosis is regulated by intrinsic circadian clocks in many species, such as mouse, rat and Sudanian grass rat (LaVail 1980; Grace et al. 1999; Bobu and Hicks 2009). However, in some species, such as in frogs and goldfish, phagocytosis peaks disappear when the ambient light is removed (Basinger & Hollyfield, 1980; Bassi & Powers, 1990). Here, to study the effect of ambient light on the phagocytosis’ rhythmicity of cone subtypes in zebrafish, the larvae were kept in constant darkness (DD) for at least 24 h before, and up until sample collection (Figure 4A). Samples were collected at the same time points and treated and analysed as previously described for the LD treated larvae. The numbers of phagosomes from UV, blue and green cone OSs were quantified with the developed analysis tool, whereas OS phagosomes from red cone OSs were calculated manually.

A two-way ANOVA analysis showed statistically significant reduction in the phagosome numbers from UV, blue and green cone OSs in DD compared to LD over 24 h (p= 5.87*10^-20, p= 5.58*10^-5 and p=7.59*10^-8, respectively) (Figure 4B). Subsequent Bonferroni Post Hoc analysis showed that the numbers of phagosomes from UV cone OSs at both peaks seen in LD (Figure 3C) were significantly lower in DD (ZT5: p= 6.58*10^-17, ZT16: p= 1.55*10^-12) (Figure 4B). Similarly, the first peak of phagosomes from blue cone OSs at ZT3 in LD was significantly dampened (p= 8.70*10^-6) under DD light condition. Additionally, the second peak of phagosomes from blue cone OSs was slightly dampened and shifted from ZT16 to ZT18. However, the difference in OS phagosome numbers between LD and DD at this new peak time point (ZT18) was not statistically significant (Figure 4B). The number of phagosomes from green cone OSs was also significantly decreased at the two time points, ZT16 and ZT23, in DD (ZT16: p=3,65*10^-8, ZT23: p=6,86*10^-5) (Figure 4B). Figure 4C summarizes how the rhythmic profiles of UV, blue and green cone OS phagocytosis alter in response to change in light condition from LD (solid lines) to DD (dashed lines). The peaks in the numbers of phagosomes from UV and blue cone OSs clearly diminish in DD light condition, but the profile of green cone phagocytosis remain more constant as clear peaks were not seen even in LD (Figure 4C).

The two-way ANOVA analysis showed that the change in light condition did not affect the total number of manually quantified phagosomes from red cone OSs throughout the 24 h (Supplementary S6). However, the Bonferroni Post Hoc test revealed a significant reduction in red cone OS phagosomes at ZT23 when LD and DD conditions were compared (p= 1.42*10^-6) (Supplementary 6). Interestingly, at ZT3, the number of phagosomes from red cone OSs was significantly higher in DD than in LD (p= 5.60*10^-4). Similar increase in OS phagosome numbers in DD at any time point were not seen with the other cone subtypes (Figure 4B).

To evaluate the contribution of UV, blue and green cone subtypes on the combined rhythmicity profile in DD condition, OS phagosome numbers for each of these cone subtypes were normalized and then summed together as described in section 3.3. The stacked bar graph containing UV, blue and green cone subtypes shows three time points during the 24 h cycle in DD condition at which the sum of the normalized values of these cone phagosomes peaked slightly (Figure 4D). Figure 4D shows that light onset-associated peak of OS phagosomes emerged already at ZT1 and continued until ZT3. This was different from the LD condition, where the peak, containing the same cone subtypes, did not emerge until ZT3 (Figure 3D). In addition, in DD, the peak after the supposed light-to-dark transition appeared at ZT18 which was later than in LD at ZT16 (Figure 3D, Figure 4D). Surprisingly, different from LD condition, a slight increase in OS numbers emerged also at ZT10 in DD. The non-normalized values of UV, blue and green cone phagosomes were then summed and averaged at each time point (Table 5). Altogether, the numbers of detected OS phagosomes over the 24 h were more constant compared to those in LD condition (Table 4, Table 5).

**Table 5.** Averages of phagosomes from UV, blue and green cone OSs per 10 µm of RPE at each time point in DD. n≥5 sections at each time point, each section represents one larva. OS: outer segment.

| Time | UV cones | Blue cones | Green cones | Average from the sum of UV, blue and green cone OSs |
| --- | --- | --- | --- | --- |
| ZT1 | 1.29 | 0.77 | 4.10 | 2.17 |
| ZT3 | 1.30 | 1.56 | 2.86 | 1.93 |
| ZT5 | 0.95 | 1.42 | 1.59 | 1.29 |
| ZT10 | 1.29 | 1.15 | 2.42 | 1.78 |
| ZT16 | 0.68 | 1.40 | 1.56 | 1.24 |
| ZT18 | 0.93 | 1.97 | 3.67 | 2.34 |
| ZT23 | 0.87 | 0.94 | 1.47 | 1.13 |
| Average of total OS phagosome count over 24 hours | 1.04 | 1.30 | 2.56 |  |

## 4. Discussion

Phagocytosis of photoreceptor outer segments has been studied intensively over several decades focusing primarily on rods. However, the rhythmic nature and regulation of the cone OS phagocytosis has remained more elusive, and to our knowledge no previous studies regarding the differences between cone subtypes have been conducted. Since cone subtypes display morphological and functional differences, divergence may also exist in the rhythmicity of the process. In addition, as previous studies have been primarily conducted with adult animals (LaVail 1980; Bobu and Hicks 2009; Ruggiero et al. 2012), rhythmicity and regulation of OS phagocytosis are largely unresolved in the developing animals. In the present study, we investigated the cyclic occurrence of OS phagocytosis of all individual cone subtypes (UV, blue, green and red cones) in cone-dominant 7 *dpf* old zebrafish larvae using immunohistochemistry, confocal microscopy and semi-automatized quantification of OS phagosomes in the RPE. We showed that all cone subtypes are phagocytosed in young, developing zebrafish and that the phagocytosis occurs at some level throughout the day. Interestingly, we found that phagocytosis of UV and blue cone OSs do exhibit rhythmicity, which was observed as two peaks in the numbers of OS phagosomes over the 24 hours. This rhythmicity dampened in the constant darkness, indicating that the process in larval zebrafish is mostly driven by the external light rather than the intrinsic circadian system.

Our data reveals that the phagocytosis of all cone subtypes is ongoing already in 7 *dpf* young zebrafish larvae (Figure 2A). As a comparison, in mice and rats, the rod OS phagocytosis process has been shown to begin around the time of eye opening: at postnatal day P14-P15 in mice and P15-P16 in rats (Lew et al. 2020). In these rod-dominant mammals, rhythmicity of the process develops and aligns with external light-cycles as well as intrinsic circadian cycles during the subsequent days as the retina and circadian systems continue to develop (Lew et al. 2020). Our data from 7 *dpf* zebrafish demonstrates two light/dark-transition-associated peaks in the combined phagosome numbers from UV, blue and green cone OSs during the 24 h circadian cycle (Figure 3D, Table 4). This result agrees with previous data combining either rods and cones or all cone subtypes but not rods and obtained from various species, such as certain mouse strains, Sudanian grass rat, chicken (*Gallus domesticus*), domestic cat, rhesus monkey (*macaca mulatta*), goldfish and zebrafish (Young 1978; O’Day and Young 1978; Anderson et al. 1980; Fisher et al. 1983; Bobu and Hicks 2009; Lewis et al. 2018; Goyal et al. 2020; Moran et al. 2022a). However, when the rhythmicity of different cone subtypes was examined separately, our quantitative data suggests that phagocytosis of zebrafish single cones (UV and blue) and the future forming double cones (green and red) are differently regulated in the early larval stages (Figure 3C, supplementary 4). We found that UV and blue cones are rhythmically phagocytosed with two daily peaks in the OS phagosome numbers. This result is supported by a previous study by Krigel et al. (2010) that utilized the neural retina leucine zipper gene knockout mouse strain (*Nrl^−/−^*) to study the rhythmicity of blue cone phagocytosis (Krigel et al. 2010). These mice completely lack rods in the retina, but the remaining photoreceptors seem to be identical to blue cones. Krigel et al. showed that in these mice, the rhythmic profile of the blue cone OS phagocytosis corresponded to the rhythm of rods in wild-type mice with a significant peak in OS phagosome numbers at 1 h after light onset and a slighter increase at 1 h after light offset. While our results revealed rhythmic nature for the phagocytosis of the UV and blue cones, the green and red cones did not show similar rhythmicity profile. We speculate this to be due to differences in the developmental phases of the cone subtypes in the larval zebrafish retina. It is commonly recognized that UV and blue cones develop and mature earlier than green and red cones (Robinson et al. 1995; Raymond et al. 1995; Schmitt and Dowling 1999; Crespo and Knust 2018). Additionally, in young zebrafish larvae, all cones appear as single cones, since the red/green double cones are visible only after 12 *dpf* (Branchek and Bremiller 1984; Allison et al. 2010; Zimmermann et al. 2018). Therefore, we assume the green and red cones to be less mature than the UV and blue cones at 7 *dpf*. More developed rhythm in UV and blue cones could also be related to the level of their activity in zebrafish at this age. As the main objective of the larvae is to survive and grow, prey capture is in the center of their behavior. In the larval zebrafish, the UV cones are known to be used primarily in prey detection (Zimmermann et al. 2018; Yoshimatsu et al. 2020), highlighting their importance for the larvae. This could partially explain the higher demand for their rhythmic and synchronized renewal already in the young individuals.

In addition to the different rhythmic profiles in the phagocytosis of the distinct cone subtype OSs, our data showed variation in the total numbers of cone-type specific phagosomes over a 24 h cycle (Figure 3C, Table 4, Supplementary S4). However, as the numbers can be affected by the used antibodies, it is not straightforward to compare the absolute numbers of OS phagosomes between the different cone subtypes. For instance, our results showing the higher number of green cone phagosomes compared to the other cone subtypes can be influenced by the used antibody. To our knowledge, an antibody specific to only green cone opsin in zebrafish is not publicly available. Therefore, we used zpr-3, an antibody that has been shown to label the OSs of green cones and rods in zebrafish (Hu et al. 2024). As fully functional rods do not appear before 15 *dpf* in zebrafish (Branchek and Bremiller 1984), it is likely that the demand for the renewal of rod precursor OS membranes is not remarkable in 7 *dpf* old larvae. However, we cannot neglect the possibility that rod precursors are renewed and phagocytosed to some extent, and this way they could affect our results showing higher numbers of quantified phagosomes labelled by zpr-3. Interestingly, our immunolabelling showed that rod opsin (O4886) and zpr-3 antibodies both in the adult and larval cryosections were similar, indicating that both antibodies label not only rods (in adults) or rod precursors (in 7 *dpf* old larvae) but also green cones in zebrafish (Supplementary S5). Therefore, we were not able to specifically label rod precursors and evaluate how much they contribute to the observed phagosome numbers in the larval cryosections. We encountered antibody specificity issues also in the labelling of red cones, which could explain the low numbers of the detected phagosomes from red cone OSs in our cryosections. We used rhodopsin [1D4] antibody (ab5417, Abcam) that has been shown to label rods in the mouse retinal samples (Ramon et al. 2007), but interestingly, Yin et al. 2012 showed that in zebrafish, this antibody labels OSs of red cones (Yin et al. 2012). In our experiments, this antibody showed noisy and unspecific labelling of the cryosections. Thus, the phagosomes from red cones were hard to distinguish from the background. We assume this to be one reason also for the failed quantification with the created semi-automatized analysis tool, as it cannot detect and segment the OS particles of red cones properly from the background. Therefore, the analysis tool should be trained more for a more reliable quantification of OS phagosomes of this cone type. Furthermore, both rhodopsin [1D4] and zpr-3 antibodies recognize and bind to an epitope at the C-terminus of the rhodopsin protein (Hu et al., 2024; MacKenzie et al., 1984, Supplementary S1). The C-terminus of the rhodopsin is digested rapidly within the RPE after OS particle engulfment, as the formed OS phagosome is fused to acidic enzymes containing endosomes (Wavre-Shapton et al. 2014). This makes the rhodopsin [1D4] and zpr-3 antibodies useful only for detecting newly engulfed OS particles. Since our UV and blue cone subtype-specific opsin antibodies bind to the more stable N-termini of their target proteins, the phagosomes both at early and later stages of digestion can be detected with these antibodies. Overall, since the used opsin-specific antibodies differ in their binding sites on the opsin proteins as well as in their specificity, it is more meaningful to compare the rhythmicity profiles than the absolute phagosome numbers between the different cone subtypes.

As our results showed rhythmicity in the phagocytosis of UV and blue cone OSs, we further studied the involvement of external light in the phagocytosis regulation. For that, we kept zebrafish larvae in constant darkness for a minimum of 24 h before sample collection, preparation and OS phagosome analyses. These constant darkness experiments showed that the impact of the external light on the cone phagocytosis was evident in the larval zebrafish with a notable decrease in the total number of OS phagosomes over a 24 h cycle when compared to the LD condition (Figure 4B, Table 4, Table 5). Moreover, in DD, the peaks in OS phagosome numbers from UV, blue and red cones diminished (Figure 4B, Figure 4C, Supplement S6). This result differs from the previous study by Krigel et al. where they showed that the rhythmicity of blue cone OS phagocytosis remained in DD condition in the *Nrl^−/−^*mouse model, and concluded that the blue cone OS phagocytosis is under circadian control (Krigel et al. 2010). On the contrary, our data indicates that in young zebrafish larvae, the rhythmicity of cone OS phagocytosis is mainly driven by the alternating light-dark cycles and not substantially regulated by the intrinsic circadian system. This finding is also supported by the study of Moran et al 2022 from 16 *dpf* larvae. Overall, circadian rhythms related to zebrafish cellular functions and behaviors show differences in the maturation rates (Reviewed by Vatine et al., 2011). It is thus plausible, that the circadian systems shown to be linked to the regulation of the OS phagocytosis in mice, rats and cultured RPE cells (Ruggiero et al. 2012; Baba et al. 2017; Milićević et al. 2021; DeVera et al. 2022), are still developing in the 7 *dpf* old zebrafish larvae. Therefore, their impact on the regulation might not be observable at that age. Nevertheless, it is worth noting that in goldfish and frogs, the OS phagocytosis peaks are primarily caused by the light-dark cycles even in the adult stages (Basinger & Hollyfield, 1980; Bassi & Powers, 1990). Zebrafish belongs to the teleost class of fishes as does goldfish. Therefore, the process could be regulated by the light cycles also in the adult zebrafish.

In conclusion, this study showed that under normal light cycle, all cone subtypes are phagocytosed in the young zebrafish larvae. Surprisingly, the rhythmic nature of the process seems to be different in the distinct cone subtypes at 7 *dpf*. For UV and blue cones, the rhythmicity is evident with two daily peaks of OS phagosomes, whereas for the red and green cone phagocytosis, clear rhythmicity was not observed. Additionally, our results revealed the influence of changes in light-dark conditions for the rhythmic bursts of phagocytosis in the 7 *dpf* old larvae, suggesting weak regulation by the intrinsic circadian clocks either related to the developmental immaturity or species-specific characteristics. Further research is needed to determine the mechanism by which the phagocytosis of the distinct cone subtype OSs is differentially regulated and whether the circadian system is involved.

## Abbreviations

RPE: Retinal pigment epithelium
OS: Outer segment
Dpf: Days post fertilization
ERG: Electroretinogram
IS: inner segment
ONL: Outer nuclear layer
OPL: Outer plexiform layer
ZT: Zeitgeber time
EM: Electron microscopy

## 5. Acknowledgements

We would like to thank Tampere Zebrafish Facility for providing and maintaining the zebrafish used in this study. We also thank Tampere Histology Facility and Tampere imaging Facility for the equipment for preparation and imaging the histological samples, respectively. We thank Milvi Ali-Alha for the help with immunolabelling. The project has been funded with support of the Silmä-ja kudospankkisäätiö (Finland), Finnish cultural foundation (Finland), Mary and Georg C. Ehrnrooth Foundation (Finland), Academy of Finland, Emil Aaltosen säätiö (Finland).

