## Supplemental materials for "Rhythmicity of photoreceptor outer segment phagocytosis differs between the cone subtypes in the larval zebrafish"

### Supplementary 1

***Table S1: Binding sites of the used opsin antibodies at their target opsin protein***

| Antibody | Labelled cells | Binding site at opsin protein, amino acids | Binding site at opsin protein, terminus |
|---|---|---|---|
| UV opsin | UV cones | 1-27 | N |
| Blue opsin | Blue cones | 1-41 | N |
| Rod opsin | Rods in various vertebrate phyla: mammal, avian, amphibian, fish (Barnstable 1980, Silver et al., 1988) | 4-10 | N |
| Rhodopsin [1D4] | Red cones in zebrafish (Yin et al., 2012) | 339-348 | C |
| zpr-3 | Green cones+Rods (Hu et al., 2024) | 320-354 | C |

### Supplementary S2

#### *A) Code S2: The full script of the semi-automatized analysis tool for quantitative analysis of outer segment (OS) phagosomes in the RPE tissue.*

```
/// Macro by Sanni Erämies
/// Version: 08.01.2025
/// Requires: MorphoLibJ, ResultsToExcel (test mode only)
/// Description:
/// This macro assists in particle analysis from RPE images using h-dome transformation
/// for peak detection. The process includes user interaction for parameter selection,
/// channel configuration, RPE area selection, and analysis based on provided parameters.
/// The results are saved as ROI files and optionally as Excel sheets for further analysis.
///
/// Key Features:
/// - User-driven RPE area selection.
/// - Automatic particle detection using  h-dome values and prominence thresholds.
/// - Outputs ROIs of detected particles and RPE area.
///
/// Sections:
/// 1. Function Definitions
/// 2. Main Macro Execution
////////////////////////////////////////////

// Function Definitions

/// Displays a dialog for user input and initializes key paths and parameters.
///
/// Inputs:
/// - None (user provides input via a dialog).
///
/// Outputs:
/// - An array containing the following:
///   - `input` (`string`): The user-selected input directory.
///   - `output` (`string`): The user-selected output directory.
///   - `h_input` (`number`): The user-defined h-value for dome detection.
function startup(){
		Dialog.createNonBlocking("RPE particle analysis");
		Dialog.addDirectory("Input Directory", getDirectory("home"));
		Dialog.addDirectory("Output Directory", getDirectory("home"));
		Dialog.addNumber("Define h-value for dome detection", 15); // Default h-value for
dome detection

		Dialog.show();
		input = Dialog.getString(); // User-selected input directory
		output = Dialog.getString(); // User-selected output directory
		h_input = Dialog.getNumber(); // User-defined h-value

		return newArray(input, output, h_input);
}

/// Allows the user to define which channels to use for analysis.
///
/// Inputs:
```

```
/// - None (user selects channels via dialog).
///
/// Outputs:
/// - An array containing the following:
///   - `rpe_ch` (`string`): The user-selected RPE channel.
///   - `particle_ch` (`string`): The user-selected particle channel.
function channelCheck(){
		Dialog.createNonBlocking("Channels");
		Dialog.addChoice("RPE:", CHANNELS, 2);
		Dialog.addChoice("Particles:", CHANNELS, 3);
		Dialog.show();

		ch1 = Dialog.getChoice(); // RPE channel
		ch2 = Dialog.getChoice(); // Particle channel

		title = "MIP";
		run("Duplicate...", "duplicate channels="+ch1+"-"+ch2+" title="+title);
		close("\\Others");
		run("Split Channels");

		rpe_ch = "C1-"+title;
		particle_ch = "C2-"+title;

		return newArray(rpe_ch, particle_ch);
}

/// Enables the user to select the RPE area for analysis. The user can refine the selection until satisfied.
///
/// Inputs:
/// - `img` (`string`): The name of the image where the RPE area will be selected.
///
/// Outputs:
/// - Adds the RPE selection to the ROI Manager with the name `rpe`.
function rpeAreaSelection(img){
		looper = true;
		setForegroundColor(255, 255, 255);
		setBackgroundColor(0, 0, 0);

		while(looper == true){
				selectWindow(img);
				run("Duplicate...", "duplicate title=RPE");

				Dialog.createNonBlocking("Info");
				Dialog.addMessage("Define RPE area to use in the analysis. \nOK when ready.");
				Dialog.show();

				run("Clear Outside");
				setAutoThreshold("Li dark");
				run("Convert to Mask");
				run("Morphological Filters", "operation=Closing element=Disk radius=10");

				run("Create Selection");
```

```
				roiManager("reset");
				roiManager("Add");
				roiManager("select", 0);
				roiManager("Rename", "rpe");
				close("RPE thresholding");

				// Show preview
				selectWindow(img);
				roiManager("show all");

				// Repeat or continue with the selection
				Dialog.createNonBlocking("Continue with the current selection?");
				Dialog.addRadioButtonGroup("Continue or re-do", 
newArray("Continue", "Restart RPE-selection") , 1, 2, "Continue");
				Dialog.show();

				buttonSelected = Dialog.getRadioButton();
				if (buttonSelected == "Restart RPE-selection") {
					close("RPE");
					close("RPE-Closing");
					roiManager("delete");
				} else {
					break;
				}
		}
}

/// Executes the h-dome detection for particle detection.
///
/// Inputs:
/// - `fname` (`string`): The filename for logging the output, used only in test mode!
/// - `image` (`string`): The image from which particles will be detected.
/// - `h` (`number`): The `h`-value for dome detection, controlling sensitivity.
/// - `sensitivity` (`number`): The sensitivity threshold for particle detection.
///
/// Outputs:
/// - Prints to the console:
///   - The number of detected particles.
///   - The area of the RPE region.
/// - Updates the ROI Manager with the detected particles.
function run_h_domes(fname, image, h, sensitivity) {
		selectWindow(image);
		mask = getTitle();
		run("8-bit");
		run("Duplicate...", "title=marker");
		run("Subtract...", "value=["+h+"]");

		run("Morphological Reconstruction", "marker=marker mask=["+mask+"] type=[By 
Dilation] connectivity=4");
		imageCalculator("Subtract create", mask,"marker-rec");
		rename("hdomes");

		run("Log");
		run("Morphological Filters", "operation=[White Top Hat] element=Disk radius=4");
```

```
			// FOR NORMAL MODE ACTIVATE
			roiManager("select", 0);
			run("Find Maxima..."); // Running this way allows preview
			roiManager("Add");
			roiManager("select", 1);
			roiManager("Rename", "detected particles");

			run("Clear Results");
			run("Set Measurements...", "mean redirect=None decimal=0");
			roiManager("Measure");
			print("---- Found Particles: "+ nResults());

			run("Clear Results");
			roiManager("select", 0);
			run("Set Measurements...", "area redirect=None decimal=0");
			roiManager("Measure");
			print("---- RPE area: " + getResult("Area", 0));
			// FOR NORMAL MODE ACTIVATE ^^^

			// FOR TEST MODE ACTIVATE
			//run("Find Maxima...", "prominence=["+sensitivity+"] output=[Point Selection]");
			//roiManager("Add");
   //roiManager("select", 1);
   //roiManager("Rename", "detected particles");

   //run("Set Measurements...", "centroid redirect=None decimal=3");
   //roiManager("select", 1);
   //roiManager("multi-measure append");

   //for (roi = 0; roi < nResults(); roi++) {
			//setResult("Sensitivity", roi, sensitivity);
			//setResult("h-value", roi, h);
			//setResult("filename", roi, fname);
   // }
   // updateResults();

   //print("---- Found Particles xy: "+ nResults);
   //run("Read and Write Excel", "no_count_column
file=["+user_input[1]+"/evaluation_measurements.xlsx] sheet=["+sheet+"] stack_results");
   //run("Clear Results");
   //roiManager("select", 0);
   // FOR TEST MODE ACTIVATE^^^
}

/// Saves detected ROIs to the output directory.
///
/// Inputs:
/// - `fname` (`string`): The filename used to save the ROIs.
/// - `dir` (`string`): The directory where the ROIs will be saved.
///
/// Outputs:
/// - Saves the ROIs with the name `fname` as a zip file in the `dir` directory.
function saveROIResultsToOutput(fname, dir){
			roiManager("Select", newArray(0)); // Select RPE and detected particles
			roiManager("Save", dir+"/"+fname+".zip");
```

```
}

/// Executes in the test mode by running h-dome detection across various parameter combinations.
///
/// Inputs:
/// - `filename` (`string`): The base filename used for results logging.
/// - `POSimage` (`string`): The image from which particles will be detected.
/// - `hArray` (`array of numbers`): An array of `h`-values for testing different sensitivities.
/// - `sensitivityArray` (`array of numbers`): An array of sensitivity thresholds to test.
///
/// Outputs:
/// - Runs `run_h_domes` multiple times with different combinations of `h` and sensitivity values.
function runTEST(filename, POSimage, hArray, sensitivityArray){
		for (i = 0; i < lengthOf(hArray); i++) {
				print("h-value: "+ hArray[i]);
				for (j = 0; j < lengthOf(sensitivityArray); j++) {
						print("sensitivity: "+ sensitivityArray[j]);
						run_h_domes(filename, POSimage, hArray[i], 
sensitivityArray[j]);
						roiManager("select", 1);
						roiManager("Delete");
						close("hdomes-White Top Hat");
						close("hdomes");
						close("marker-rec");
						close("marker");
				}
		}
		print("DONE, all saved");
}

// Main Macro Execution

testMode = false; // Set to true for testing
sensitivity = newArray(5, 10, 20, 40, 60); // Sensitivity thresholds for tests
hvalues = newArray(5, 15, 50, 100, 300); // h-dome values for tests
CHANNELS = newArray("1", "2", "3", "4"); // Available channels

user_input = startup();
filelist = getFileList(user_input[0]);

for (i = 0; i < lengthOf(filelist); i++) {
		if (endsWith(filelist[i], ".nd2")) {
				print("Processing: "+filelist[i]);
				run("Bio-Formats Importer", "open=[" + user_input[0] + filelist[i] + 
"] autoscale color_mode=Default rois_import=[ROI manager] view=Hyperstack 
stack_order=XYCZT");
				filename = File.getNameWithoutExtension(getTitle());
				getDimensions(width, height, channels, slices, frames);
				run("Z Project...", "projection=[Max Intensity]");
				run("Gaussian Blur...", "sigma=1 stack");
				Stack.setDisplayMode("composite");

				analysis_channels = channelCheck();
				rpe = analysis_channels[0];
				particle = analysis_channels[1];
```

```
			rpeAreaSelection(rpe);

			if(testMode == true){
				runTEST(File.getNameWithoutExtension(filelist[i]), 
particle, hvalues, sensitivity);

		saveROIResultsToOutput(File.getNameWithoutExtension(filelist[i]), user_input[1]);
			} else {
				run_h_domes("", particle, user_input[2], 50);
				selectWindow(particle);
				roiManager("select", 1);
				waitForUser("OK to save results");
				saveROIResultsToOutput(filename, user_input[1]);
				close("*");
			}
		}
}
```

**Supplementary S2**

***B) Table S2: Parameters and their definitions and mathematical equations needed for the evaluation of the performance of the analysis tool's peak detection algorithm***

| **Parameter** | **Definition** | **Equation** |
|---|---|---|
| Precision | The proportion of detected peaks that were correct | $Precision = \frac{TP}{(TP + FP)}$ |
| Ground truth | The manually annotated local intensity maxima | |
| Sensitivity (Recall) | The proportion of ground truth peaks that were successfully detected | $Sensitivity = \frac{TP}{(TP + FN)}$ |
| F1-score (F1) | The harmonic mean of precision and sensitivity, providing a single measure of performance | $F1 = \frac{2x(Precision\ x\ sensitivity)}{(Precision + sensitivity)}$ |
| False Positive Rate (FPR) | The proportion of negative ground truth peaks incorrectly classified as positive | $FPR = \frac{FP}{FP + TP}$ |
| True positive (TP) | A detected peak that was within three pixels of a ground truth peak | |
| False positive (FP) | A detected peak that could not be matched to any ground truth peak | |
| False Negative (FN) | A ground truth peak that was not matched to any detected peak | |

### Supplementary S2

## C)

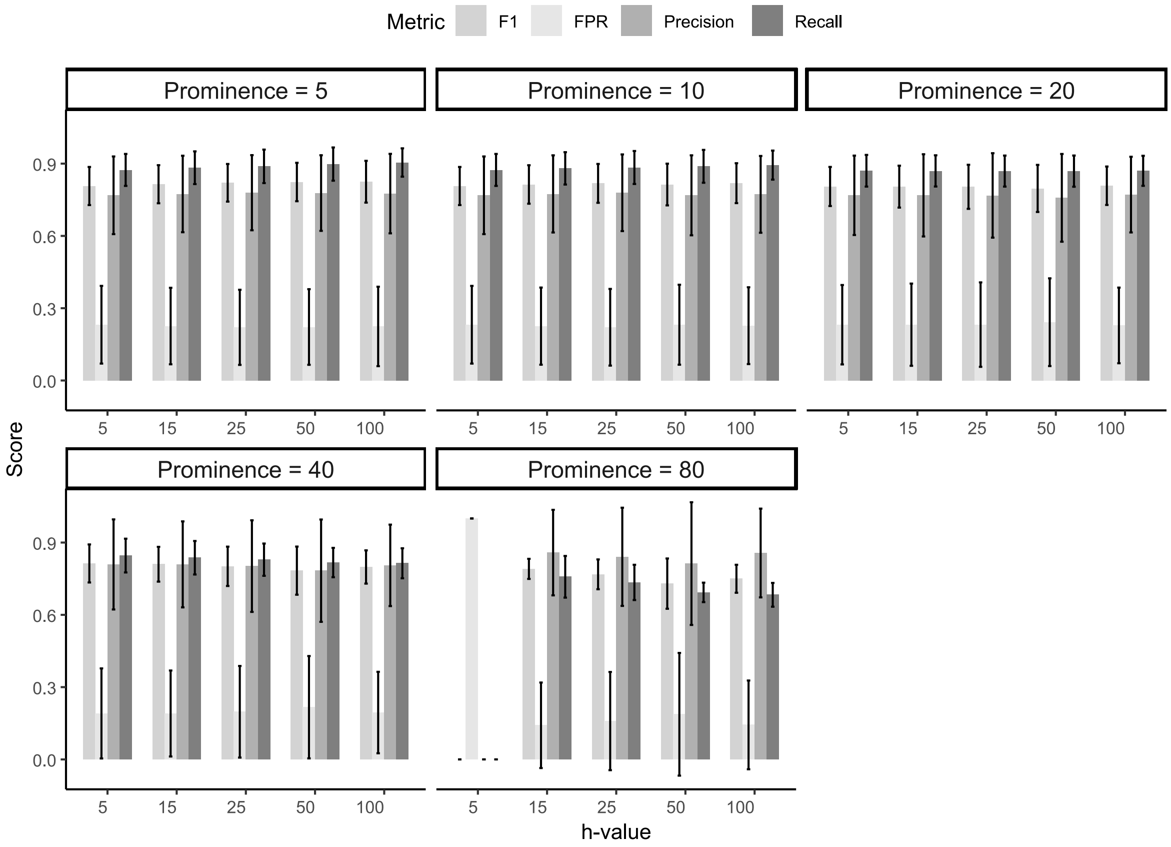


***Fig. S2 Comparison of the performance of the analysis tool's peak detection algorithm using different combinations of five h-dome values and five prominence thresholds across three randomly selected sample images.** The plots show the calculated score values for precision, Recall (sensitivity), F1-score (F1), and false positive rate (FPR) on the Y-axis for each h-dome-prominence threshold combination, allowing for an evaluation of the algorithm's overall detection reliability. The tested h-values are on the X-axis. The results suggest that the h-value has minimal impact on the algorithm's performance to detect True positive (TP) intensity peaks. Moreover, the data indicates that the performance remains nearly identical with the prominence threshold values set between 5 and 20 with each h-values. However, around prominence threshold value 80, and especially in combination with low h-values, the algorithm's performance seems to decline. This is likely due to over-filtering. For the final analysis tool, h-value was set to 15 and the prominence threshold values is defined by the experimenter specifically for each image sample.*

**Supplementary S3**

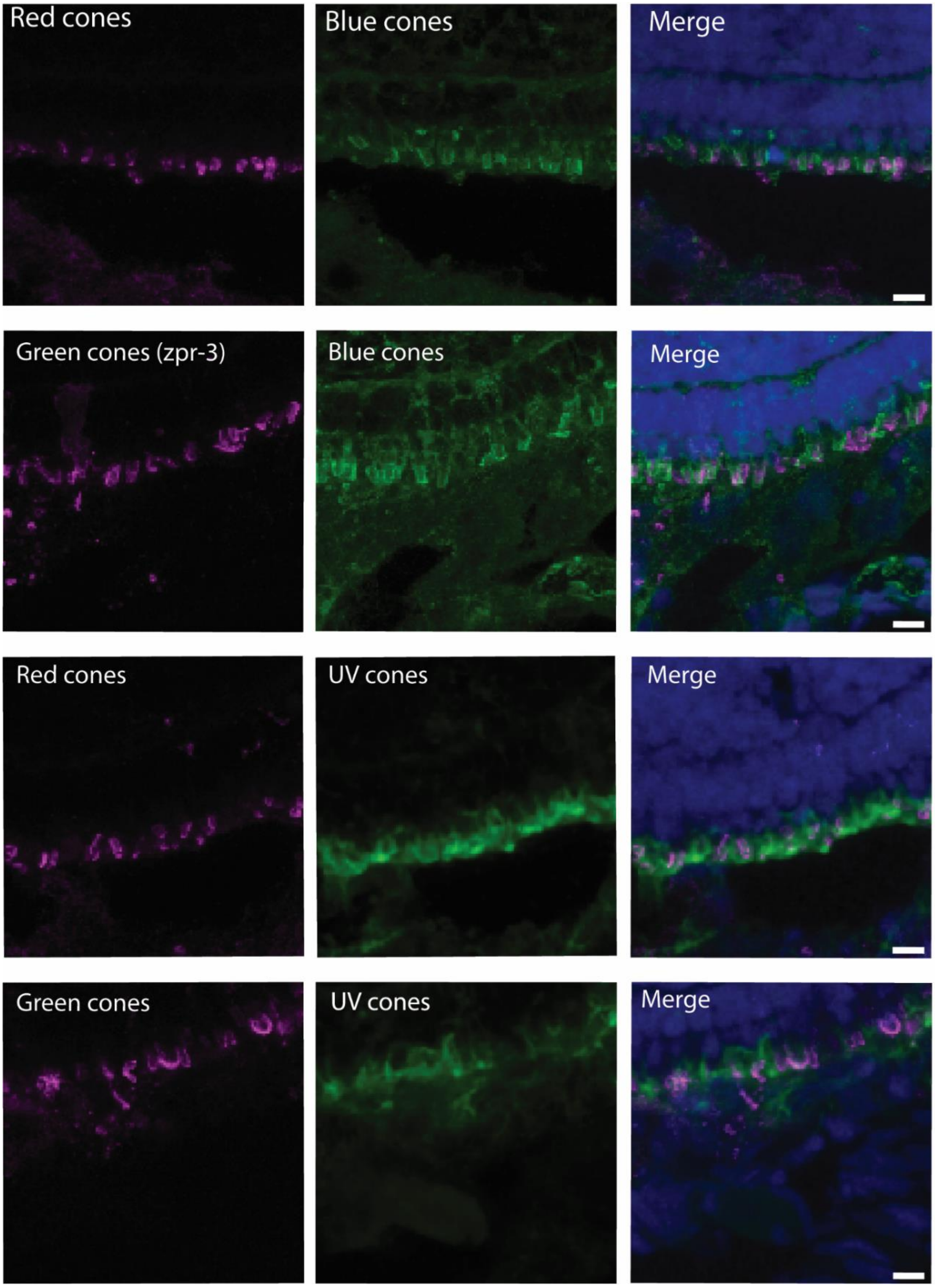


**Fig S3. Immunofluorescence labelling of 7 *dpf* old zebrafish cone subtypes.** *Different antibody combinations (magenta and green) were used to label simultaneously two different cone subtypes together with DAPI (Blue). The confocal images of all the combinations show that the OSs of different cone subtypes appear in the same layer in zebrafish at 7 dpf. Scale bars 5 µm. OSs: outer segments*

**Supplementary S4**

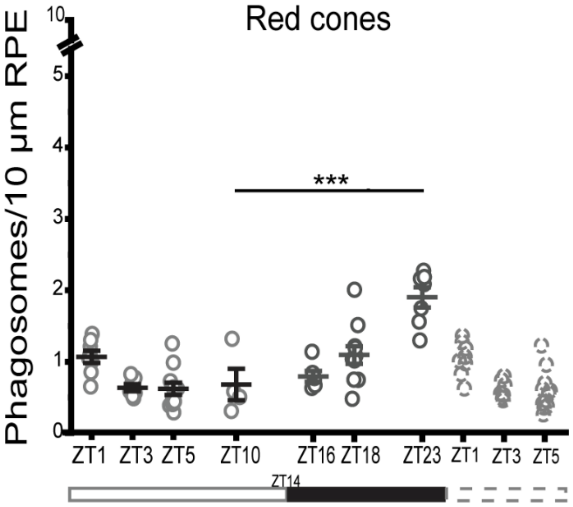


| Time | Red cones in LD |
|---|---|
| ZT1 | 1.07 |
| ZT3 | 0.63 |
| ZT5 | 0.62 |
| ZT10 | 0.69 |
| ZT16 | 0.82 |
| ZT18 | 1.12 |
| ZT23 | 1.93 |
| Average of total OS phagosome count over 24 hours | 0.98 |

***Fig S4. The numbers of phagosomes from red cone OSs in the RPE over the 24 h period in normal light-dark cycle.*** *The scatter plot shows the number of phagosomes from red cone OSs per 10 µm of RPE at each studied time point. Data is represented as individual samples (circles) together with the mean line ± SEM. One-way ANOVA analysis was used to show statistically significant differences in OS phagosome numbers over the 24 hours. Subsequent Bonferroni's post hoc test revealed significant increase in OS phagosomes at ZT23 (1.93 phagosomes/10 µm RPE) compared to the baseline at ZT10 (0.69 phagosomes/10 µm RPE) (***p<.001). The white bar and the black bar represent the light and dark periods of the day, respectively. The dashed bar shows again the three first time points of the day. The table shows averages of the phagosomes from the red cone OSs per 10 µm of RPE at each time point under normal light-dark cycle. Phagosomes were quantified from the length of the entire RPE tissue in each whole eye section. N≥5 sections at each time point, each section represents one larva. RPE: Retinal pigment epithelium, ZT: Zeitgeber time, White bar under the graph: light period of the day, Black bar under the graph: dark period of the day, OS: outer segment.*

### Supplementary S5

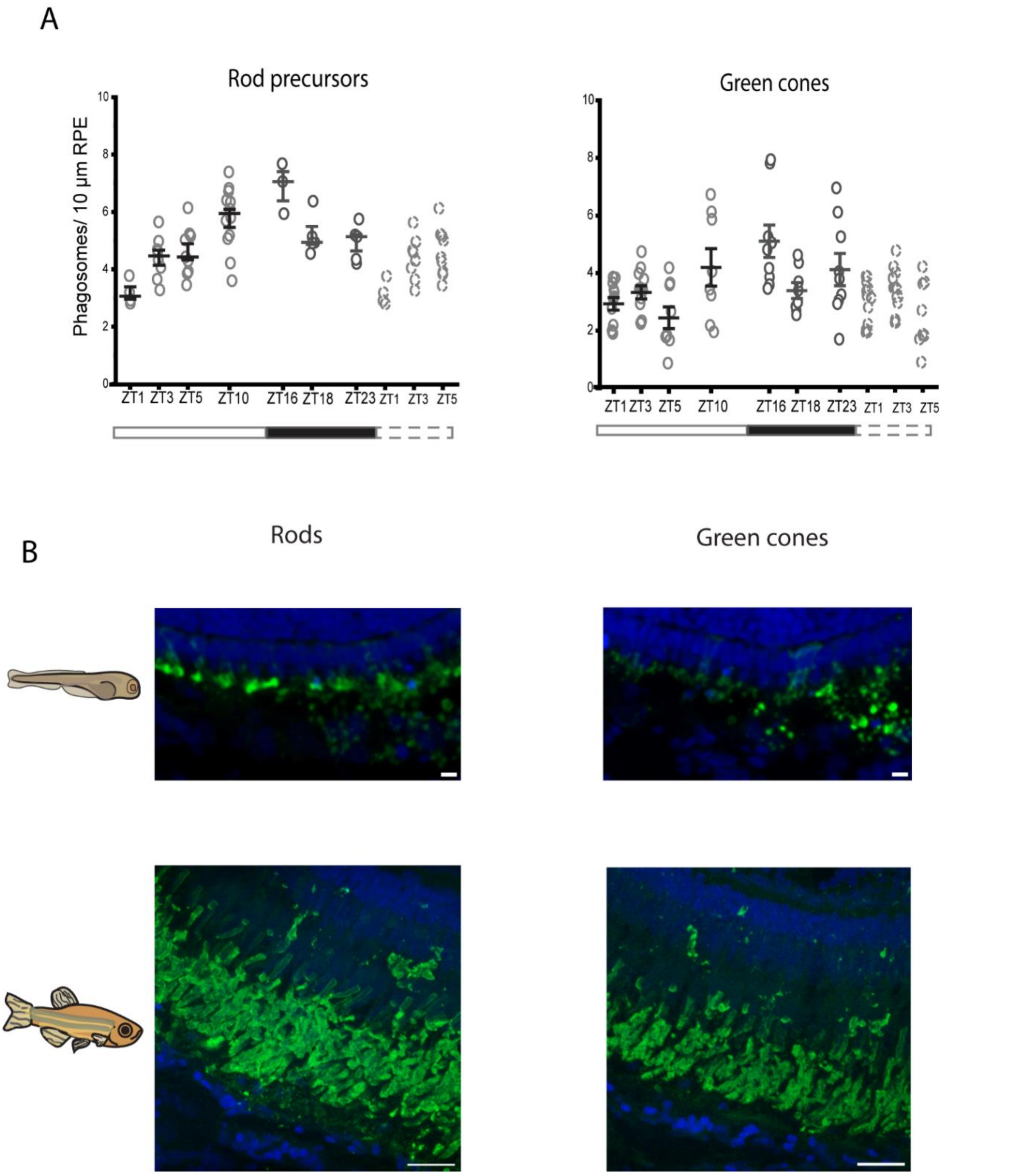


***Fig S5. Quantitative data and image data of rods and green cones*** *A) Graphs show the number of phagosomes from rod precursor OSs and green cone OSs per 10 µm of RPE at each studied time point. Data is represented as individual samples (circles) together with the mean line ± SEM. Rhythmicity of phagocytosis of rod precursor OSs shows similar trend to that of green cone OSs. The white bar and the black bar represent the light and dark periods of the day, respectively. The dashed bar shows again the three first time points of the day. B) Immunofluorescent labelling of 7 dpf larval (wild-type) and adult (Fli-eGFP strain) zebrafish cryosections using DAPI stain (Blue) together with either rod opsin antibody (O4886) or zpr-3 to visualize rods (green) and green cones (green), respectively. In adult sections (lower row), these antibodies seem to label both rod and cone layers. Scalebar in larval sections: 5 µm and in adult sections: 20 µm.*

### Supplementary S6

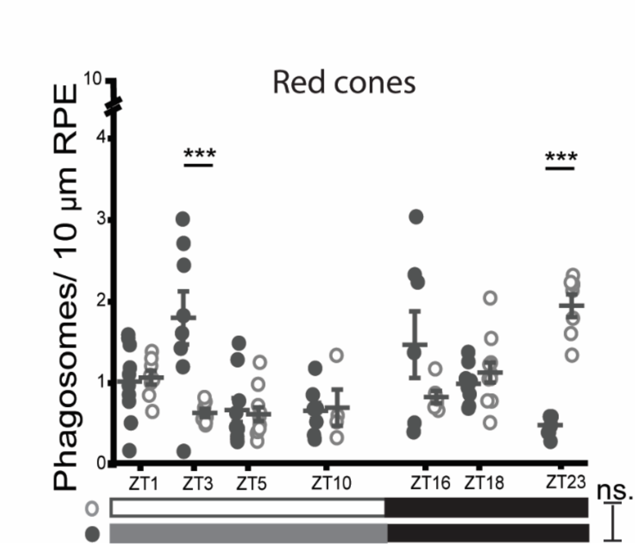


| Time | Red cones in DD |
|---|---|
| ZT1 | 1.02 |
| ZT3 | 1.80 |
| ZT5 | 0.67 |
| ZT10 | 0.65 |
| ZT16 | 1.46 |
| ZT18 | 0.98 |
| ZT23 | 0.47 |
| Average of total OS phagosome count over 24 hours | 0.99 |

**Fig S6. *The numbers of phagosomes from red cone OSs in the RPE* over the 24 hours in constant darkness.**

The *scatter plot shows the numbers of phagosomes from red cone OSs per 10 µm of RPE at each studied time point in normal light cycle (LD, white) and in constant darkness (DD, Dark grey). Data is represented together with the mean line ± SEM. At ZT23, the peak seen in LD was diminished in DD (***p<.001), while a clear peak emerged at ZT3 (***p<.001). Two-way ANOVA analysis was used to show that there was non-significant (ns.) difference in the numbers of phagosomes from red cone OSs over the 24 hours between LD and DD conditions. The table shows averages of the phagosomes from red cone OSs per 10 µm of RPE at each time point under constant darkness. Phagosomes were quantified from the length of the entire RPE tissue in each whole eye section. N≥5 sections at each time point, each section represents one larva. RPE: Retinal pigment epithelium, ZT: Zeitgeber time, OS: outer segment.*
